# DNA-PK enforces the S/G2 checkpoint in ATM-deficient cells to limit ATR inhibitor efficacy

**DOI:** 10.64898/2026.10.25.761677

**Authors:** Timothy B. Branigan, Roshen Alharthi, Bose S. Kochupurakkal, Jie Hao, Steffie Revia, Joachim Albers, Mu-Yan Cai, Yuqing Jiao, Kelsey McQueen, Sirisha Mukkavalli, Ian A. Roundtree, Gregory M. Cote, Prafulla C. Gokhale, Kalindi Parmar, Alan D. D’Andrea, Astrid Zimmermann, Geoffrey I. Shapiro

**Author notes:** **Corresponding Author:** Geoffrey I. Shapiro, MD, PhD, Dana-Farber Cancer Institute, 450 Brookline Avenue, Mayer 446, Boston, MA 02215.

## Abstract

ATR inhibition has been proposed as a strategy for tumors under replication stress or those harboring ATM deficiency. However, anti-tumor activity is limited by DNA-PK and ATM compensation. Here, we have delineated distinct roles for DNA-PK and ATM during ATR inhibitor exposure. Both kinases are activated in a cell cycle-dependent manner at the S/G2 boundary following ATR inhibition. DNA-PK imposes the S/G2 checkpoint and promotes completion of DNA replication, whereas ATM detects persistent aberrant DNA replication structures with associated DNA damage in late S and G2/M to establish a G2/M checkpoint promoting DNA repair. In ATM-deficient cells and patient tumors, DNA-PK is activated following ATR inhibition and assumes greater versatility by not only enforcing the S/G2 checkpoint but also by detecting aberrant replication structures in S phase in place of ATM, allowing cells to tolerate ATR inhibition. Consequently, DNA-PK inhibition synergizes with ATR inhibition in ATM-deficient cells and enhances ATR inhibitor efficacy in ATM-deficient PDX models. These data support clinical evaluation of combined ATR and DNA-PK inhibition in ATM-deficient cancers.

## Introduction

Ataxia telangiectasia and Rad3-related (ATR) is a critical kinase enabling cancer cells to tolerate replication stress. The presence of single-stranded DNA replication structures, a biochemical consequence of replication stress, activates ATR to coordinate the completion of DNA replication with cell cycle progression^1, 2^. Oncogenic alterations that induce accelerated DNA replication create a dependency on ATR to enforce an S/G2 checkpoint that allows for the detection and repair of problematic DNA replication structures^1, 3^. To establish the S/G2 checkpoint, ATR engages CHK1 to limit CDK1 activation and thereby prevent premature initiation of the mitotic cell cycle program via a positive-feedback loop between cyclin/CDK1 and the MMB-FOXM1 transcriptional complex^4, 5^. ATR also is activated by R-loops at centromeres during mitosis and functions to activate Aurora B via CHK1, thereby establishing the spindle assembly checkpoint that times the M/G1 transition^6^.

In cells under replication stress that harbor aberrant single-stranded DNA replication structures, ATR inhibition is expected to impair their resolution, further exacerbating replication stress, and to facilitate mitotic entry prior to the completion of DNA replication, resulting in tumor cell lethality^7^. Abrogation of the S/G2 checkpoint is required for ATR inhibitor efficacy as knockout of the MMB-FOXM1 components, LIN54 or FOXM1, prevents activation of the mitotic program and has been shown to attenuate the loss of ATR-CHK1 signaling^4, 8^.

Unresolved problematic DNA replication structures may be processed into double-strand DNA breaks that can be detected by the DNA damage response (DDR) regulator ATM^9, 10^. ATM is recruited to double-strand DNA breaks, where it detects DNA damage by depositing γ-H2AX, an epigenetic marker of stressed chromatin^2, 11^. ATM also promotes DNA end resection by phosphorylating the MRE11-RAD50-NBS1 (MRN) complex that allows for RAD51-mediated repair via homologous recombination (HR), and limits cell cycle progression by phosphorylating CHK2 to establish a G2/M checkpoint^2, 12, 13, 14^. Consistent with these ATM functions, ATM activity promoted formation of RAD51 foci following ATR inhibition in gastric cancer cells indicating that ATM was mobilizing the DDR response to repair DNA damage^15^. In chronic lymphocytic leukemia cells subjected to ATR inhibition, depleting ATM increased origin firing and the abundance of phospho-histone H3 (pH3)-positive mitotic cells, leading to the accumulation of under-replicated DNA^16^. These results suggest that ATM can directly offset absent ATR function. Indeed, loss of ATM increases ATR inhibitor sensitivity in multiple experimental systems^15, 16, 17^, leading to the hypothesis that ATM deficiency could serve as a predictive biomarker for ATR inhibitor response^18, 19^. However, in clinical trials, ATR inhibitors as monotherapy have demonstrated only modest response rates in ATM-deficient tumors^20, 21, 22^.

Although ATR inhibition activates ATM, the extent to which ATM signaling compensates for the loss of ATR signaling may be highly context dependent. In ATM WT U2OS osteosarcoma cells, DNA-PK, another sensor of DNA damage, rather than ATM, activated a CHK1-dependent checkpoint cells that limited origin firing following ATR inhibition^23^. In contrast, both ATM-CHK2 and DNA-PK-CHK1 pathways were activated by ATR inhibition in pancreatic cancer cells, with DNA-PK capable of activating CHK2 in the absence of ATM signaling^17^. However, only combined ATR and ATM inhibition, and not combined ATR and DNA-PK inhibition, was synergistic in the ATM-proficient cell lines tested.

The extent to which DNA-PK or ATM individually compensate for loss of ATR signaling by restricting premature activation of the mitotic program to reestablish the S2/G2 checkpoint, or by mobilizing a DNA damage response to resolve persisting single-stranded DNA replication structures, remains to be elucidated. In this study, we sought to establish a conceptual framework to understand the interplay between DNA-PK and ATM and the checkpoints they activate following ATR inhibition in both ATM-proficient and -deficient cells. We report that DNA-PK and ATM are activated following ATR inhibition to respectively promote the S/G2 and G2/M cell cycle checkpoints in response to increased CDK1, as well as CDK2 activity. DNA-PK antagonizes the S/G2 transition following ATR inhibition and promotes DNA replication, whereas ATM functions after DNA-PK activity to detect DNA damage in late S or G2/M and mobilizes the DDR response in G2/M. In ATM-deficient cells, DNA-PK plays multifaceted roles following ATR inhibition, and not only enforces an S/G2 checkpoint but also detects DNA damage in Late S phase in place of ATM. Consistent with multiple functions of DNA-PK during ATR inhibitor exposure in ATM-deficient cells, DNA-PK inhibition enhanced ATR inhibitor efficacy in several ATM-deficient tumor models. These data support distinct roles for ATM and DNA-PK in compensating for impaired ATR signaling and point to combined ATR and DNA-PK inhibition as a treatment strategy for ATM-deficient tumors, explaining the limited efficacy of ATR inhibitor monotherapy in this setting.

## Results

### Deregulation of CDK1/2 after ATR inhibition activates DNA-PK and ATM to prevent mitotic progression at the S/G2 boundary

To assess how DNA-PK and ATM compensate for a loss of ATR signaling, we investigated their activation in SNU719 gastric cancer cells following treatment with the ATR inhibitor tuvusertib (ATRi)^24^. DNA-PK and ATM were both activated, evidenced by phosphorylation at their respective activation sites, pDNA-PK (S2056) and pATM (S1981), during exposure to a concentration of 500 nM ATRi over 24 hours, which was also sufficient to induce replication stress markers, including pS4/S8 RPA32, pS33 RPA32^25^ and pS824 KAP1 (pKAP1)^4, 26^, as well as γ-H2AX (**Fig. 1a**). We next determined if activation of DNA-PK and ATM coincided with the establishment of checkpoint control to detect aberrant DNA replication structures and slow cell cycle progression, allowing for the resolution of these structures through DNA replication and repair. Levels of pDNA-PK and pATM, as well as activated pS345 CHK1 (pCHK1) and γ-H2AX, were monitored by flow cytometry over a 24-hour time course of ATRi treatment for the accumulation of cellular populations with high levels of these markers (**Supplementary Fig. 1a, b**; **Fig. 1b-e**). Levels of activated pT68 CHK2 (pCHK2) were also monitored by immunoblot (**Supplementary Fig. 1c**). While activation of CHK1 is initially limited by ATR inhibition, cells with high levels of pCHK1 and γ-H2AX were induced at 12 hours and increased at 16 hours during ATRi exposure (**Supplementary Fig. 1a; Fig. 1b, c**). Similarly, levels of pCHK2 increased at 12 hours and peaked at 16 hours (**Supplementary Fig. 1c**). Coinciding with these events, populations of pDNA-PK^High^ and pATM^High^ cells increased modestly at 12 hours, and both populations were significantly induced after 16 hours of ATRi treatment (**Supplementary Fig. 1b**; **Fig. 1d, e**). These results indicate that activation of DNA-PK and ATM following ATR inhibition corresponds with CHK1/2 activation and establishment of cell cycle checkpoint control, evidenced by the accumulation of cells at the interface between Late S and G2/M populations between 12-16 hours post ATRi (**Supplementary Fig. 1d, e**; **Fig. 1f, g**). Activation of DNA-PK and ATM also occurs at the S/G2 boundary as the pDNA-PK^High^ and pATM^High^ cells are induced at the same interface (**Fig. 1f, h, i**; **Supplementary Fig. 1f, g**) further supporting the idea that DNA-PK and ATM are activated in response to ATR inhibition to enforce a checkpoint at the S/G2 boundary.

**Fig. 1.**
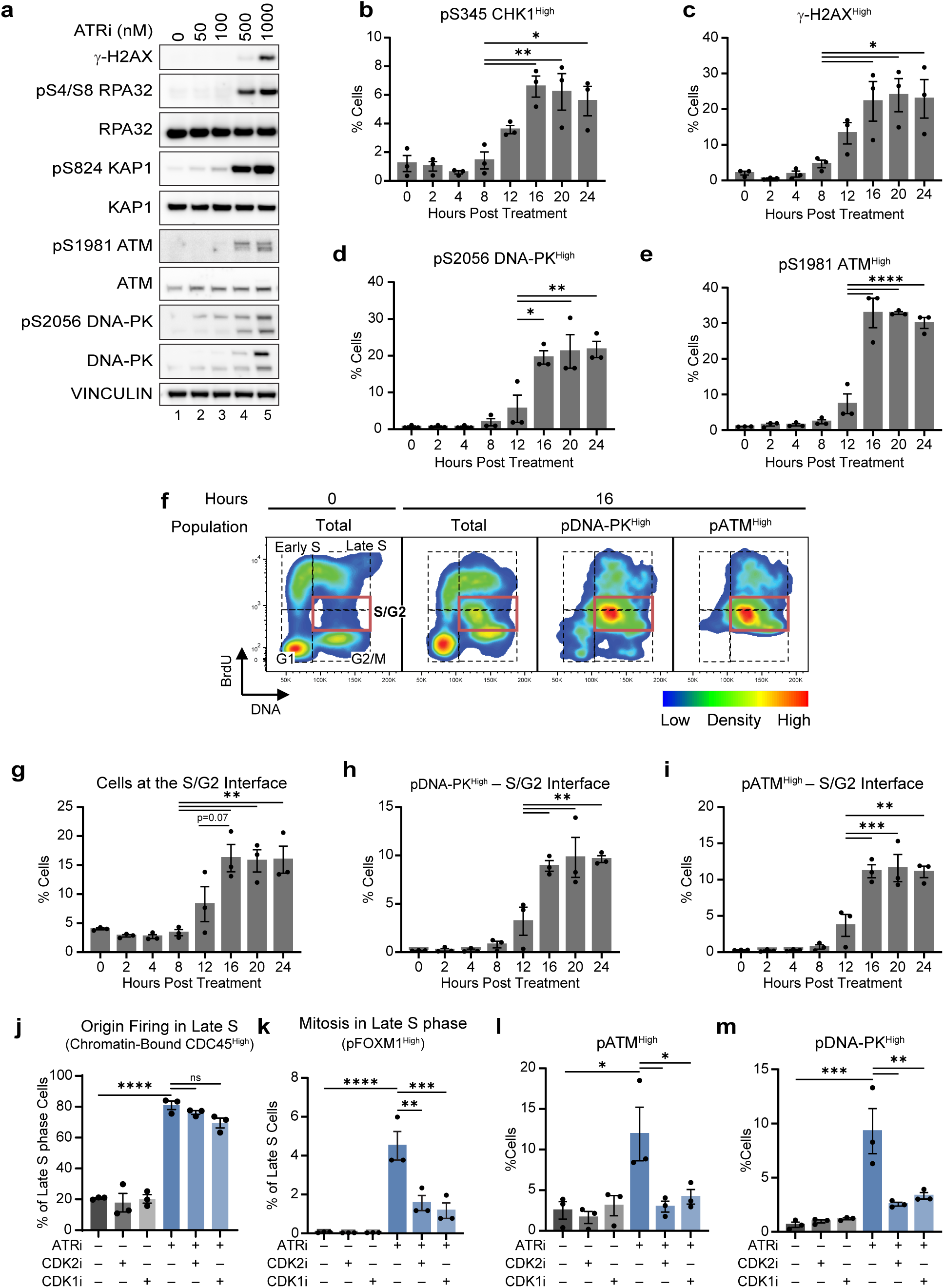
DNA-PK and ATM are activated following ATR inhibition at the S/G2 boundary to premature activation of the mitotic program. **a** Immunoblot showing activation of DNA-PK and ATM after ATRi. SNU719 cells were treated with denoted concentrations of tuvusertib (ATRi) for 24 hours. Whole cell extracts were generated and probed with the denoted antibodies. **b-i** DNA-PK and ATM are activated at an S/G2 checkpoint following ATR inhibition. SNU719 cells were treated with 500 nM ATRi for the denoted time points and analyzed by flow cytometry for γ-H2AX and pCHK1 levels or pDNA-PK and pATM, BrdU incorporation and DNA content, N=3**. b** % of cells that are pS345 CHK1High (pCHK1High), **c** % of cells that are γ-H2AXHigh, **d** % of cells that are pS025 DNA-PKHigh (pDNA-PKHigh), **e** % of cells that are pS1981 ATMHigh (pATMHigh ), **f** Cell cycle distribution of samples at 0 (left) and 16 hours (middle left) post ATRi and cell cycle distribution of the pATMHigh (middle right) and pDNA-PKHigh (right) cells at 16 hours post ATRi. S/G2 interface denoted by red box, **g** % of cells at the S/G2 interface, **h** % of cells that pDNA-PKHigh at the S/G2 interface, **i** % of cells that pATMHigh at the S/G2 interface. **j-m** Premature mitosis after ATR inhibition required for DNA-PK and ATM activation. SNU719 cells were treated with 500 nM ATRi ±10 μM CDK2i and/or 2.5 μM CDK1i for 16 hours and levels of chromatin-bound CDC45, EdU, and DNA content were analyzed by chromatin flow cytometry, N=3, or levels of γ-H2AX and pFOXM1 or pDNA-PK, BrdU incorporation, and DNA content were analyzed by flow cytometry, N=3. % of Late S phase cells that are CDC45High. **k** % of Late S phase cells that are pFOXM1High **l** % of cells that are pATMHigh **m** % of cells that are pDNA-PKHigh.* p<0.05. ** p<0.01. *** p<0.001.

During the S/G2 transition, the ATR-CHK1 pathway restricts CDK1 activity to allow for the completion of DNA replication before activation of the mitotic cellular program^4, 5, 27^. The ATR-CHK1 pathway also has been implicated in limiting origin firing during early S phase^28, 29, 30^. Loss of ATR-CHK1 signaling is therefore expected to promote the mitotic program, as well as induce excessive origin firing in a CDK-dependent manner. It is not clear whether subsequent replication fork collapse and DNA damage occurs via nucleotide and RPA exhaustion or via mitotic levels of CDK1 activity that also trigger replication fork disassembly^31^. Therefore, it is important to distinguish whether DNA-PK and ATM are activated following ATR inhibition to limit origin firing or to prevent premature mitosis at the S/G2 boundary.

To determine whether activation of DNA-PK and ATM at the S/G2 boundary following ATR inhibition are driven by excessive origin firing or premature mitosis, DNA-PK and ATM activation was assessed following ATR inhibition in the presence of a CDK1 inhibitor at sufficient concentrations to prevent premature S to G2 transition but not excessive origin firing. CDC45 loading is the rate limiting step for origin firing^32^ and levels of chromatin-bound CDC45 increased in S phase cells following ATR inhibition (**Supplementary Fig. 2a-c**). Levels of CDK1 activity can be monitored by phosphorylation of the mitotic transcription factor FOXM1 at T600 (pFOXM1)^4, 5^. High levels of pFOXM1 are typically restricted to G2/M (**Supplementary Fig. 2d**) but ATR inhibition induced high levels or pFOXM1 in Late S phase (**Supplementary Fig. 2e**; **Fig. 1j**). The submaximal concentration of the CDK1i inhibitor RO3306 that blocks premature S/G2 transition but not DNA replication following inhibition of ATR-CHK1 pathway^4, 5^, blocked the induction of pFOXM1^High^ cells in Late S phase following ATR inhibition but failed to restrict high levels of CDC45 loading (**Fig. 1j, k**). CDK2 inhibition, with the CDK2-specific inhibitor CDK2-in-4, also restricted the induction of pFOXM1^High^ cells in Late S phase but not excessive CDC45 loading following ATRi inhibition indicating that CDK2 inhibition can also be used to separate excessive origin firing and a premature mitotic program following ATR inhibition. Importantly, the induction of pDNA-PK^High^ and pATM^High^ cells following ATR inhibition was blocked by the inclusion of submaximal CDK1i or CDK2i (**Fig. 1l, m**). This indicates that excessive origin firing is not sufficient to initiate DNA-PK and ATM-mediated compensation following ATR inhibition. Taken together, these results indicate that premature CDK1/2 activity after ATR inhibition triggers activation of DNA-PK and ATM at the S/G2 interface to limit premature mitosis at the S/G2 boundary.

We also studied the induction of γ-H2AX following ATR inhibition. In this context, γ-H2AX marks problematic DNA replication structures that persist in the presence of high CDK1 activity that undergo controlled breakage to facilitate their resolution^33^. Importantly, γ-H2AX induction after ATR inhibition preferentially occurred in cells with high levels of the CDK1 substrate pT600 FOXM1 (**Supplementary Fig. 2f**) and submaximal CDK1i or CDK2i were sufficient to limit the accumulation of γ-H2AX^High^ cells after ATR inhibition (**Supplementary Fig. 2h**). Therefore, a CDK-mediated premature mitotic program is required for the accumulation of problematic DNA lesions following ATR inhibition, supporting the idea that DNA-PK and ATM activation in response to ATR inhibition mitigates this premature mitosis.

### DNA-PK and ATM establish distinct cell cycle checkpoints to compensate for ATR inhibition that correspond with the detection and resolution of aberrant DNA replication structures and associated DNA damage

While DNA-PK and ATM are both activated following ATR inhibition, ATM has been identified as the dominant compensator in multiple cell lines^15, 16, 17^ and DNA-PK in other cell lines^17, 23^ leaving the interplay between these kinases following ATR inhibition incompletely understood. To determine the contribution of DNA-PK and ATM activities in the establishment of cell cycle checkpoints after ATR inhibition, ATR-inhibited gastric cancer cells were co-treated with a DNA-PK inhibitor, peposertib (DNA-PKi)^34^ or an ATM inhibitor, lartesertib (ATMi)^35^. Levels of pCHK1, pCHK2 and γ-H2AX were assessed by flow cytometry or immunoblot. The induction of pCHK1 following ATRi is prevented by inhibition of either DNA-PK or ATM (**Fig. 2a**) whereas pCHK2 induction after ATR inhibition was only blocked by ATM inhibition (**Fig. 2b**, lanes 5 and 6), suggesting that DNA-PK and ATM have separable roles in establishing an S/G2 or a later G2/M checkpoint following ATR inhibition. Notably, levels of pCHK2 increased after combined ATR and DNA-PK inhibition (**Fig. 2b**, lanes 5 and 7). These results indicate that although DNA-PK and ATM are both activated at the S/G2 interface after ATRi exposure, they compensate for ATR inhibition sequentially. DNA-PK activity is rate-limiting prior to ATM activity, as ATM was able to compensate for the loss of both DNA-PK and ATR signaling, further supporting the possibility that DNA-PK and ATM establish temporally distinct checkpoints following ATR inhibition.

**Fig. 2.**
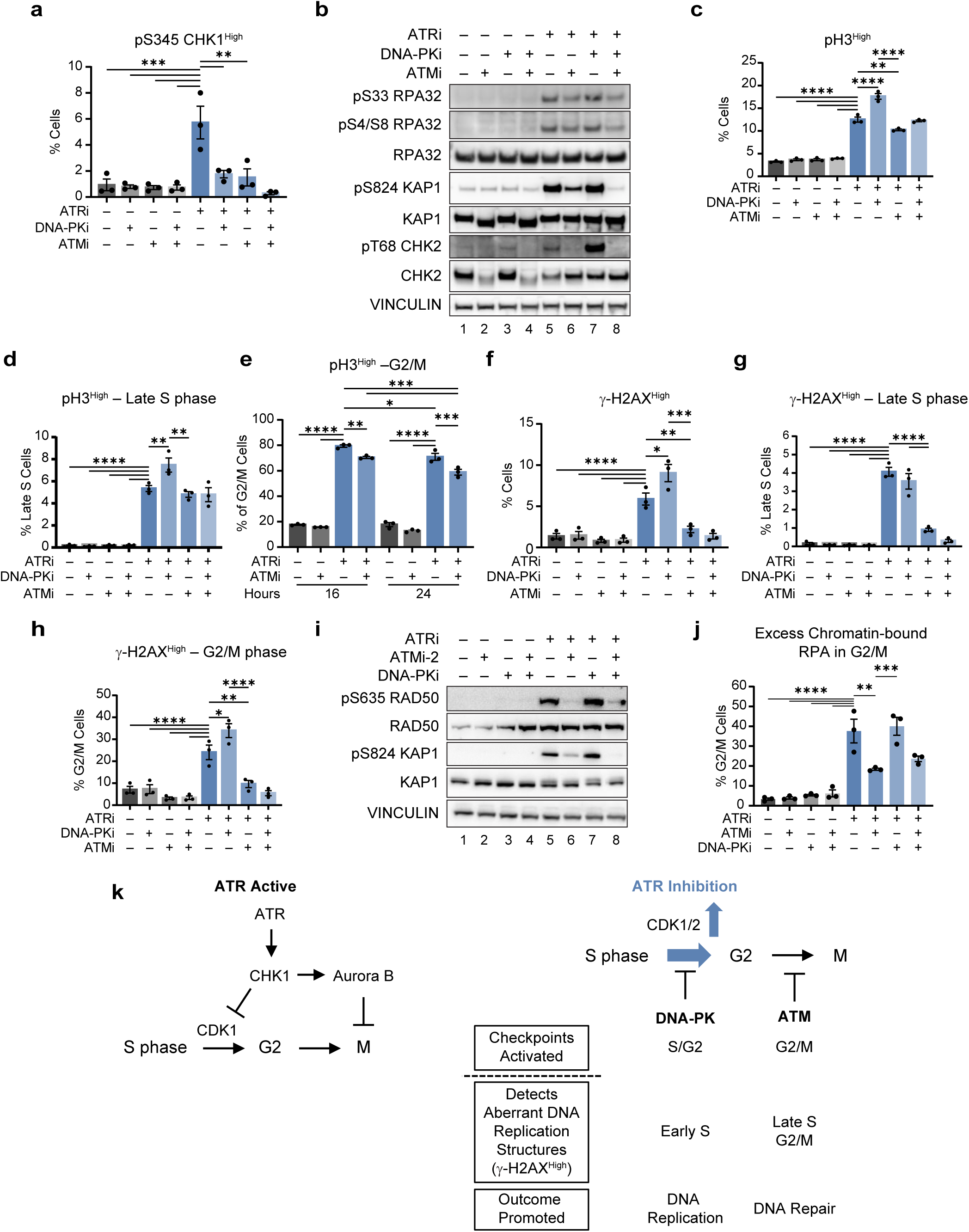
Distinct roles for DNA-PK and ATM in compensating for ATR inhibition. **a** DNA-PK and ATM activate CHK1 following ATRi. % of cells that are pCHK1High after treating SNU719 cells with 500 nM ATRi (tuvusertib) 2.5 μM DNA-PK (Peposertib) and/or ± 2.5 μM ATMi (lartesertib) for 24 hours, N=3. **b** ATM activates CHK2 following ATRi. Immunoblot of pCHK2 and DNA replication stress marker levels after treating SNU179 cells with 500 nM ATRi ± 2.5 μM ATMi and/or 2.5 μM DNA-PK for 16 hours. Whole cell extracts generated and probed with denoted antibodies. **c-h** Differing roles of DNA-PK and ATM in enforcing cell cycle checkpoints and detecting DNA damage. SNU719 cells were treated with 500 nM ATRi ± 2.5 μM DNA-PKi and/or 2.5 μM ATMi for 16 hours or 16 and 24 hours for (**e**). Levels of γ-H2AX, pH3, EdU incorporation, and DNA content were analyzed by flow cytometry, N=3. **c** % of cells that are pH3^High^, **d** % of Late S phase cells that are pH3^High^, **e** % of G2/M cells that are pH3^High^ at 16 and 24 hours post treatment following ATRi ± ATMi, **f** % of cells that are γ-H2AX^High^, **g** % of Late S phase cells that are γ-H2AX^High^, **h** % of G2/M cells that are γ-H2AX^High^. **i** Immunoblot of pRAD50 and pKAP1 induction in SNU719 after treating SNU179 cells with 500 nM ATRi ± 2.5 μM ATMi-2 and/or 2.5 μM DNA-PK for 24 hours. Whole cell extracts generated and probed with denoted antibodies. **j** % of G2/M cells with Excess RPA. SNU719 cells were treated with 500 nM ATRi ± 2.5 μM ATMi, and/or 2.5 μM DNA-PK for 16 hours. Chromatin flow cytometry was performed for chromatin-bound RPA levels, EdU incorporation, and DNA content, N=3. **k** Diagram of cell cycle regulation by ATR (left) and of ATM-DNA-PK compensation network after ATR inhibition on (right). * p<0.05. ** p<0.01. *** p<0.001. **** p<0.0001.

To further investigate the establishment of the S/G2 and G2/M checkpoints by DNA-PK or ATM following ATR inhibition, levels of the mitotic marker pH3 were assessed throughout cell division (**Supplementary Fig. 3a**). High levels of pH3 were induced by ATR inhibition in Late S phase, in addition to G2/M, leading to an increase in cells in a mitotic state (**Supplementary Fig. 3b**; **Fig. 2c-e**). Co-treatment of ATR-inhibited cells with DNA-PKi increased the levels of pH3^High^ cells in Late S phase (**Fig. 2d**), indicating that DNA-PK activity following ATR inhibition enforces the S/G2 checkpoint prior to the induction of pH3. In contrast, ATMi reduced the levels of pH3^High^ cells overall, particularly in G2/M but not late S phase (**Fig. 2d, e; Supplementary Fig. 3c, d**) as ATRi exposure persisted from 16 to 24 hours. ATM inhibition did not prevent the accumulation of pH3^High^ cells at 24 hours follow ATR inhibition when nocodazole was added at 12 hours post treatment, while cyclin B1 levels were significantly decreased in samples co-treated with ATR and ATM inhibitors compared to vehicle (**Supplementary Fig. 3e, f**) suggesting that ATR and ATM work together to maintain the G2/M state in mitotic cells. Similar to ATR, ATM promotes the spindle assembly checkpoint in mitosis^36^ suggesting that the loss of pH3 and in ability to maintain cyclin B1 levels in G2/M following combined ATR and ATM inhibition reflects premature progression through mitosis and on to the next G1. Consequently, these results indicate that ATM activity establishes a G2/M checkpoint, allowing pH3 to accumulate at G2/M.

The aberrant replication structures resulting from premature cell cycle progression after ATR inhibition are typically processed to double-strand breaks that are marked with γ-H2AX^7, 37^. The detection of problematic DNA replication structures following ATR inhibition, denoted by induction of γ-H2AX^High^ cells, was largely dependent on ATM activity (**Fig. 2f**). The accumulation of γ-H2AX^High^ cells was assessed throughout cell division by flow cytometry to determine if the detection of these aberrant structures coincided with the S/G2 or G2/M checkpoints and were DNA-PK or ATM-dependent. ATM activity was required for the detection of these structures following ATR inhibition during Late S phase and G2/M but not during the early cell cycle (**Fig. 2f, g**; **Supplementary Fig. 3g, h**). Although induction of γ-H2AX^High^ cells is predominantly dependent on ATM, there was also some dependence on DNA-PK since co-application of DNA-PKi with ATRi reduced the detection of γ-H2AX^High^ cells during Early S phase. Notably, this decrease in γ-H2AX^High^ cells following combined ATR and DNA-PK inhibition was accompanied by a corresponding ATM-dependent increase in G2/M (**Fig. 2h**) further supporting the idea that DNA-PK-mediated compensation for ATR inhibition occurs at an earlier point in cell division than ATM-mediated compensation.

Consequently, ATM activity promoted the DDR response in G2/M following ATR inhibition as the ATM inhibitor KU55933 (ATMi-2) blocked the ATRi-induced phosphorylation of the MRN complex component, RAD50 (**Fig. 2i**). The MRN complex performs DNA end resection in G2 needed for homologous recombination and the restart of stalled replication forks, resulting in the accumulation of RPA on chromatin to protect the ssDNA ends of the resected DNA strands^13, 38, 39^. Inhibition of ATM activity following ATRi treatment blocked the induction of G2/M cells with excess chromatin-bound RPA (**Fig. 2j**; **Supplementary Fig. 3i**), suggesting that ATM promotes end resection as part of the DDR response in G2/M to compensate for the loss of ATR signaling.

Together, these data indicate that ATM and DNA-PK have distinct roles in compensating for ATR inhibition (**Fig. 2k**). DNA-PK functions before ATM by responding to elevated CDK1/2 activity in S phase to enforce the S/G2 checkpoint thereby promoting the completion of DNA replication. ATM functions later in Late S and G2/M to detect aberrant DNA replication structures and to establish a G2/M checkpoint that allows for mobilization of the DDR response to facilitate the resolution of these persisting structures and associated damage. The sequential nature of DNA-PK and ATM in compensating for ATR inhibition explains the dominance of ATM in many cells, while the distinct roles for DNA-PK and ATM raise the potential for DNA-PK activity to become rate limiting in the absence of ATM.

### DNA-PK compensates for ATR inhibition in ATM-deficient cells

Given that ATM compensates for ATR inhibition after DNA-PK, we asked if compensation for ATR inhibition occurred in ATM-deficient cells. Notably, the ATRi sensitivity of SNU719 ATM KO was decreased compared to combined ATR and acute ATM inhibition (ATMi) in parental SNU719 cells (**Supplementary Fig. 4a**) raising the possibility that compensation pathways for ATR inhibition could be active in ATM-deficient cells. The replication stress markers, pKAP1 and pRPA, as well as γ-H2AX, were still induced by ATRi in ATM KO SNU719 cells and SNU1 gastric cancer cells that harbor *ATM* frameshift mutation and lack ATM expression (**Fig. 3a**). Further, pCHK1 and pCHK2 were induced in SNU1 cells after 8 hours of ATRi treatment as pKAP1, pRPA, and γ-H2AX were upregulated (**Supplementary Fig. 4b**) indicating that ATM-deficient cells activate a replication stress response in response to ATR inhibition. In both ATM KO SNU719 and SNU1 cells, the induction of these markers coincided with activation of DNA-PK (**Fig. 3a**; **Supplementary Fig. 4b**), suggesting that DNA-PK could be driving compensation for ATR inhibition in ATM-deficient cells, and account for the modest activity of ATRi monotherapy in the treatment of ATM-deficient cancers.

**Fig. 3.**
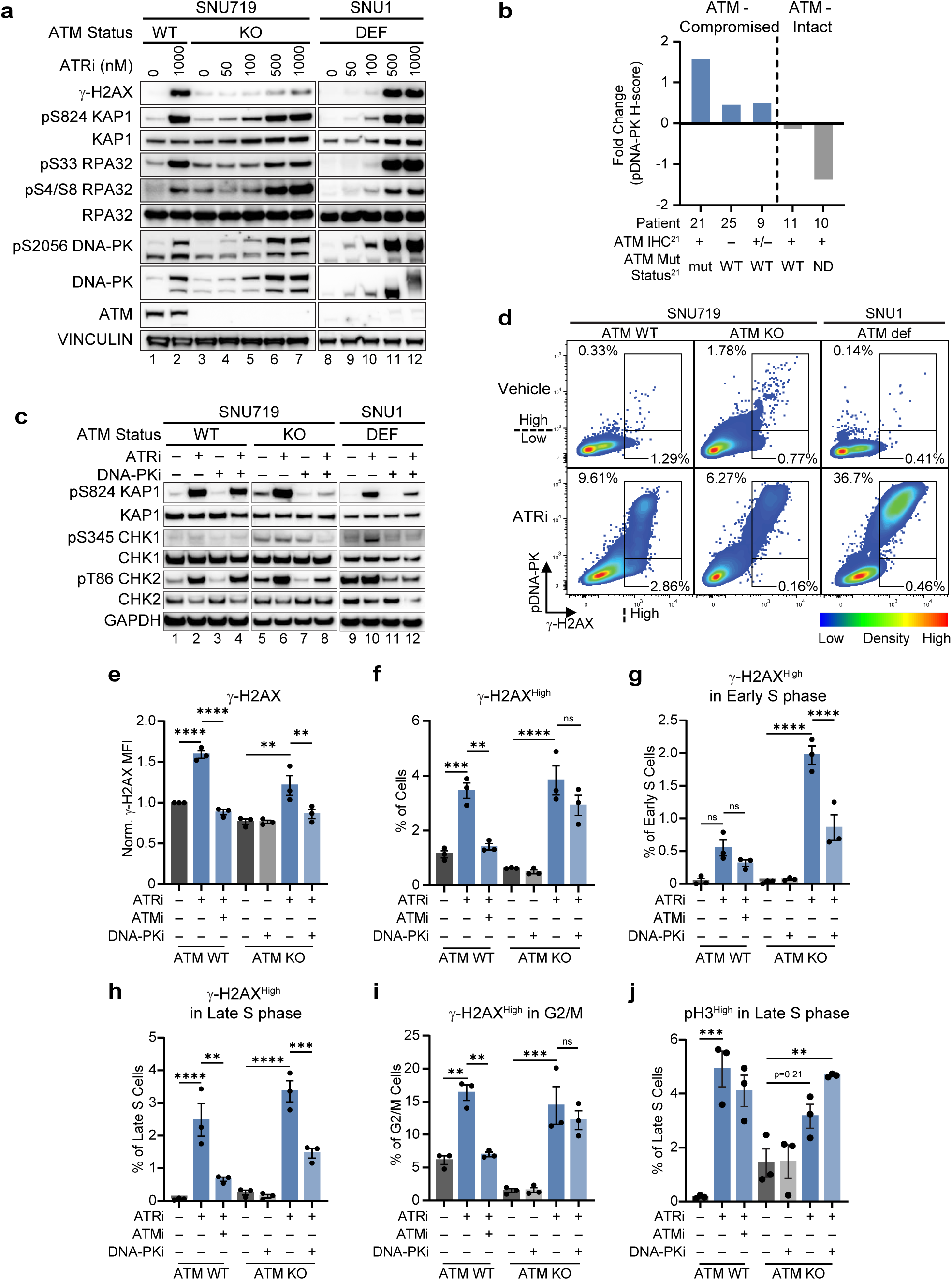
DNA-PK compensates for ATR inhibition in ATM-deficient cells. **a** Induction of RS markers in ATM-deficient cells. SNU1, ATM WT, and ATM KO SNU719 cells were treated with the denoted concentrations of ATRi (tuvusertib) for 24 hours. Whole cell extracts were generated and were probed with the denoted antibodies. **b** Fold change in pDNA-PK levels from pre-treatment sample to on-treatment sample in patients treated with ATRi-2 (berzosertib). ATM IHC positivity and mutation status listed below patient. **c** DNA-PK-dependent activation of CHK1/2 in ATM-deficient cells. SNU1, ATM WT, and ATM KO SNU719 cells were treated with 500 nM ATRi ± 2.5 μM DNA-PKi for 16 hours. Whole cell extracts were generated and were probed with the denoted antibodies. **d** γ -H2AX^High^ levels only in occur in pDNA-PK^High^ cells following ATRi in ATM-deficient cells. Plot comparing pDNA-PK and γ-H2AX levels with the pDNA-^PKHigh,^ γ -H2AX^High^ and pDNA-PK^Low^, γ -H2AX^High^ gated and % populations denoted. SNU1, ATM WT, and ATM KO SNU719 cells were treated with 500 nM ATRi for 16 hours analyzed for pDNA-PK and γ-H2AX by flow cytometry. **e-l** DNA-PK detects DNA damage in S phase and reinforces the S/G2 checkpoint following ATRi in ATM deficient cells. ATM WT and ATM KO SNU719 cells were treated with 500 nM ATRi, 2.5 μM ATMi, and/or 2.5 μM DNA-PK as denoted for 16 hours. Levels of γ-H2AX, pH3, EdU incorporation, and DNA content were analyzed by flow cytometry, N=3. **e** Mean Fluorescence Intensity of γ-H2AX, **f** % of cells that are γ-H2AX^High^, **g** % of Early S phase cells that are γ-H2AX^High^, **h** % of Late S phase cells that are γ-H2AX^High^, **i** % of G2/M cells that are γ-H2AX^High^, **j** % of Late S phase cells that are pH3^High^. * p<0.05. ** p<0.01. *** p<0.001. **** p<0.0001.

To assess if DNA-PK activation occurred in patients with ATM-deficient tumors treated with an ATRi, levels of pDNA-PK were assessed by IHC in 5 pre- and on-treatment tumor biopsy pairs procured during a Phase 2 clinical trial of the ATRi berzosertib (NCT03718091)^21^. ATM was compromised in tumors from 3 of these patients (either *ATM*-mutant or ATM-heterogeneous/negative by IHC) while 2 of the patients had tumors with ATM intact (*ATM* WT or ATM-positive by IHC). All 5 patients were on study for a comparable period before experiencing progressive disease, indicating that ATM deficiency did not afford improved clinical benefit^21^. Importantly, levels of pDNA-PK increased following berzosertib treatment in tumors harboring *ATM* mutation or low expression, but not in tumors that were *ATM* wild-type with positive expression by IHC (**Fig. 3b**; **Supplementary Fig. 4c, d**) indicating that DNA-PK was activated in response to ATR inhibition in ATM-deficient tumors.

To determine whether DNA-PK activation established cell cycle checkpoint control in ATM-deficient cells following ATR inhibition, we assessed activation of CHK1 and CHK2 following ATR inhibition in ATM KO SNU719 and SNU1 cells. In contrast to parental SNU719 cells, DNA-PK activity was required for CHK2 activation following ATR inhibition in ATM-deficient cells (**Fig. 3c**). DNA-PK-dependent CHK1 activation also increased following ATR inhibition in the ATM-deficient cell lines as pS345 CHK1 was induced in a DNA-PK-dependent manner, indicating that DNA-PK activates checkpoint control following ATR inhibition in ATM-deficient cells. Notably, the induction of the replication stress marker pKAP1 following ATR inhibition, typically associated with ATM activity, was also DNA-PK-dependent in ATM-deficient cells (**Fig. 3c**), further suggesting that DNA-PK adopts the compensatory role of ATM after ATR inhibition. In ATM WT SNU719, γ-H2AX^High^ cells were induced in both pDNA-PK^High^ and pDNA-PK^Low^ populations (**Fig. 3d**), whereas in ATM KO SNU719 and SNU1 cells, γ-H2AX^High^ cells were only induced in pDNA-PK^High^ cells, suggesting that DNA-PK is detecting aberrant DNA replication structures and associated DNA damage following ATR inhibition in the absence of ATM. DNA-PK activity was indeed required for maximal induction of γ-H2AX in ATM KO SNU719 cells after ATRi (**Fig. 3e**). However, a γ-H2AX^High^ population was still induced in the ATRi + DNA-PKi-treated cells (**Fig. 3f**), albeit with diminished levels of γ-H2AX (**Supplementary Fig. 5a**), highlighting the potential for additional compensatory pathways following ATRi that promote the resolution of replication stress. DNA-PK activity was required for the detection of aberrant replication structures and associated DNA damage in Early and Late S phase but not G2/M (**Fig. 3g-i**; **Supplementary Fig. 5b**) potentially explaining the presence of γ-H2AX^High^ cells despite DNA-PK inhibition. The induction of a G2/M population of γ-H2AX^High^ cells in the ATM KO cells following ATRi + DNA-PKi was unexpected but PLK3 and PKM2 have been reported to directly phosphorylate S139 on H2AX in response to stress or DNA damage^40, 41^, while MST1 will phosphorylate H2AX to promote apoptosis^42^, suggesting alternative compensatory or death pathways could be activated. Additionally, in ATM KO cells following ATR inhibition, high levels of γ-H2AX are induced in both pFOXM1^low^ and pFOXM1^High^ populations (**Supplementary Fig. 5c**) and more pDNA-PK^High^ cells are induced in Early and Late S phase (**Supplementary Fig. 5d**). Therefore, DNA-PK activity in ATM-deficient cells following ATR inhibition leads to the accumulation of γ-H2AX^High^ cells during Early and Late S phase, in contrast to ATM WT cells, where aberrant DNA replication structures and associated DNA damage are detected in Late S and G2/M.

Combined ATRi + DNA-PKi significantly increased the levels of pH3^High^ cells over ATRi monotherapy alone in ATM KO cells (**Supplementary Fig. S5e**). While there was a modest increase of pH3^High^ cells in G2/M in ATM KO cells following combined ATRi + DNA-PKi treatment (**Supplementary Fig. S5f**), ATRi + DNA-PKi but not ATRi monotherapy significantly induced a pH3^High^ cell population in late S phase (**Fig. 3j**) indicating that DNA-PK inhibition was needed in addition to ATR inhibition to compromise the S/G2 checkpoint in ATM KO cells. Together, these data indicate that DNA-PK is activated by ATR inhibition in ATM-deficient backgrounds to detect aberrant replication structures with associated DNA damage, to promote DNA replication in damaged cells, and to enforce the S/G2 checkpoint.

### DNA-PKi enhances ATRi efficacy in ATM-deficient tumor models

To determine if DNA-PK compensatory activity limited ATRi efficacy in an ATM-deficient background, we next treated SNU719 ATM WT and KO cells with the ATR inhibitor gartisertib (ATRi-3)^43^ and DNA-PKi (peposertib). The combination of ATRi-3 and DNA-PKi was synergistic in the ATM KO cells at lower concentrations than ATM WT cells (**Fig. 4a-d**). DNA-PK inhibition also increased the efficacy of ATRi-3 in SNU719 ATM KO but not ATM WT xenografts (**Supplementary Fig. 6a-c**). Combined ATRi (tuvusertib) + DNA-PKi therapy was also tested in 4 ATM-deficient, *KRAS*-mutant PDX models derived from patients with gastric cancer, colorectal cancer, or non-small cell lung cancer (**Fig. 4e**; **Supplementary Fig. 6d**). DNA-PKi enhanced the efficacy of ATRi in the GAPF158, REX037, LFXA-2184, and LFXA-2155 models (**Fig. 4f-4i**), converting modest tumor growth inhibition to tumor regression. These data indicate that DNA-PK inhibition enhances ATRi efficacy in a variety of ATM-deficient tumors.

**Fig. 4.**
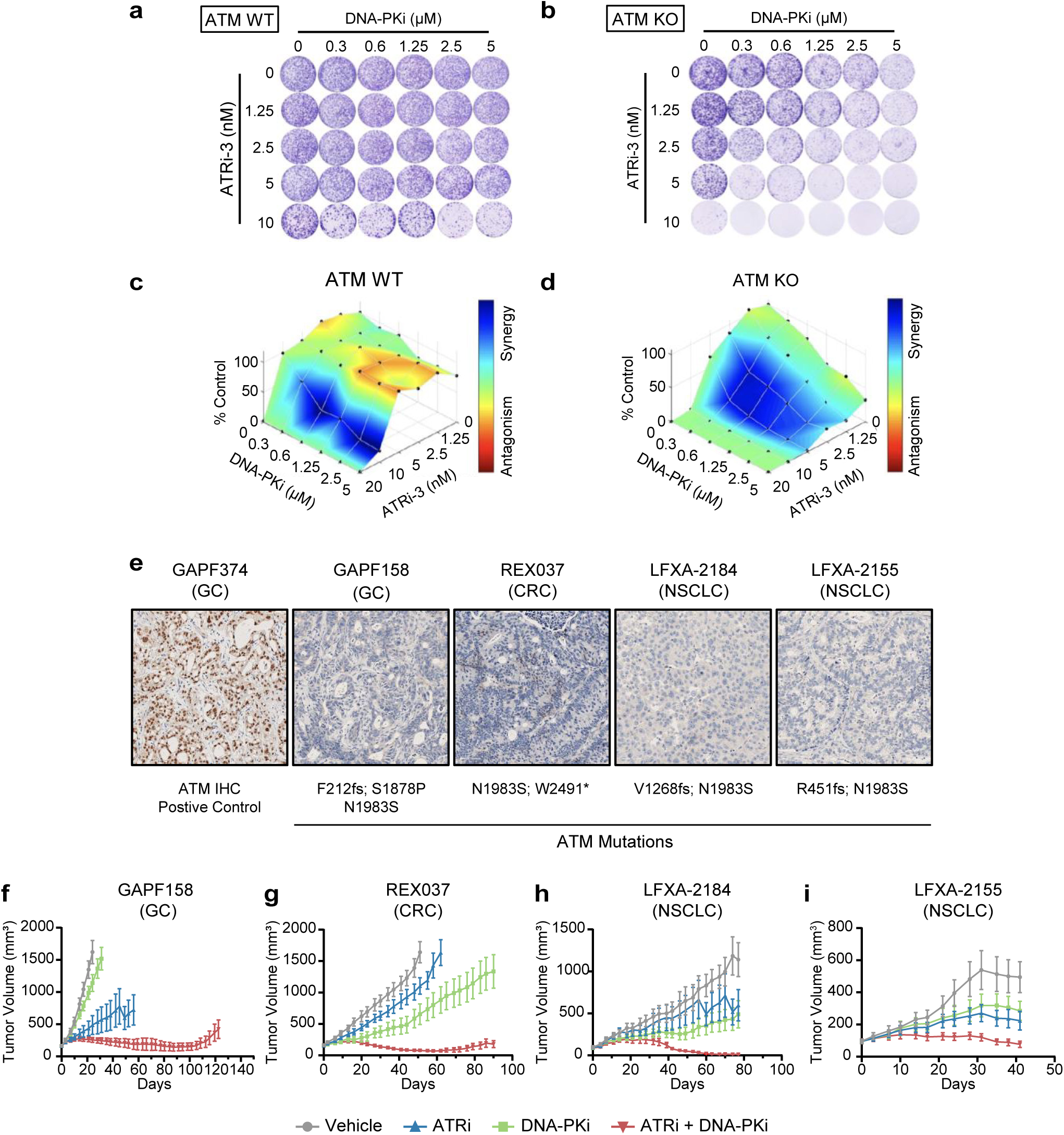
DNA-PK inhibition enhances ATR inhibitor efficacy in ATM-deficient tumors. **a-d** Synergy between ATR and DNA-PK inhibition in ATM KO SNU719 cells. **a** and **b** Representative colony formation assays in **a** ATM WT and **b** ATM KO SNU719 cells at concentrations of ATRi-3 (gartisertib) and DNA-PKi (peposertib) denoted. **c** and **d** Bliss synergy levels between ATRi-3 and DNA-PKi in **c** ATM WT and **d** ATM KO cells on the dose-experimental combination dose-response surface from colony formation assays in (**a**) and (**b**) respectively. **e-j** Efficacy studies of DNA-PK and ATR inhibition combination therapy in the ATM-deficient, KRASmut GFP158, RWX037, LFXA-2154, and LFXA-2155 PDX tumor models. **e** Immunohistochemical staining of ATM in PDX with listed ATM mutations. Tumor volumes for the ATM-deficient **f** GFP158 (ATRi (tuvusertib); dose: 25 mg/kg), **g** REX037 (25 mg/kg), **h** LFXA-2184 (10 mg/kg), and **i** LFXA-2155 (10 mg/kg) PDX models following tuvusertib (model-dependent; QD) + DNA-PKi (peposertib) (100 mg/kg; BID) combination therapy.

## Discussion

In this study, we developed a cell cycle-dependent framework for understanding how DNA-PK and ATM compensate for ATR inhibition. We determined that DNA-PK and ATM compensate for the loss of ATR signaling by sequentially establishing distinct cell cycle checkpoints at the S/G2 boundary, involving sequential S/G2 and G2/M checkpoints, respectively, providing a way to reconcile prior studies that have differed on which kinase constitutes the dominant compensation pathway. The observation that ATM functions in the Late S and G2/M phases versus DNA-PK earlier in S phase explains the ATM dominance observed in ATR-inhibited gastric and pancreatic cancer cells, where ATM is activated after combined ATR and DNA-PK inhibition. However, the ability of DNA-PK activity to enforce the S/G2 checkpoint independent of ATM explains the modest response rates of ATR inhibitor monotherapy in ATM-deficient tumors.

The observation that DNA-PK and ATM are activated at the S/G2 boundary seemingly diverges from prior observations that implicate the ATR-CHK1 pathway in regulating early S phase but highlights the interconnected molecular architecture of cell division. S phase, G2, M are often thought of as discrete periods of time but, in actuality, they reflect different states/points on a continuous spectrum of kinase activity centered on CDK1/2^44, 45^. Consequently, when DNA replication is prevented by CDC7 inhibition + CDC6 degradation, cells do not persist in an S phase state as they exit G1 but instead proceed onto mitosis from G1 with mitotic kinase activation^46^. Thus, the ATR-CHK1 pathway, activated by DNA replication, plays a critical role in restricting the cell to the S phase portion of the CDK1/2 spectrum where DNA replication can occur without impediment from the mitotic cellular program^5^. Inhibiting the ATR-CHK1 pathway allows cell cycle positive feedback loops to prematurely increase CDK activity in S phase resulting in an increase in Early S phase events^29, 30^, such as origin firing, which require a lower level of CDK activity while sufficient CDK activity accumulates to eventually trigger premature activation of the mitotic cellular program^1, 4^.

In this study, we utilized concentrations of CDK1i and CDK2i that allowed us to differentiate cell cycle events that require high vs. low CDK activities. The 2.5 µM concentration of CDK1i allows for a pool of CDK activity that does not origin firing but is sufficient to prevent premature mitotic entry. Blocking early S phase events, as previously observed, requires higher doses of CDK1i (i.e., 5 μM)^29^ that also restrict the positive feedback loops that drive premature mitotic entry^4^. Using low concentration of CDK1i, we showed that high levels of origin firing are not sufficient for DNA-PK/ATM activation after ATR inhibition, whereas premature activation of the mitotic program is required. Although not sufficient, high levels of origin firing are likely also required for DNA-PK and ATM activation, as excessive origin firing generates the DNA replication structures that become problematic with premature mitotic entry. Together, the juxtaposition of this study with prior reports points to two necessary steps in replication catastrophe following ATR inhibition: (1) the generation of an excess of DNA replication structures from high origin firing; and (2) the inability to resolve these replication structures prior to the progression from S phase into G2. The results generated here demonstrate that it is this second step that triggers compensation by DNA-PK and ATM following ATR inhibition.

Activation of DNA-PK and ATM did not occur immediately after ATR inhibition but was evident by 12-16 hours. To increase the CDK1/2 activity in S phase that is needed for DNA-PK and ATM activation, the cell must express cyclins A2 and B1 at mitotic levels prior the completion of S phase, requiring an intact MMB-FOXM1-CDK1 positive feedback loop to drive *CCNA2* and *CCNB1* expression^4, 5^. Consistent with the need for accumulating these cyclins, antagonizing the premature activation of FOXM1 transcription factor in Late S phase through submaximal CDK1i treatment is sufficient to prevent the activation of ATM/DNA-PK and the induction of stressed chromatin. As the *CCNA2* and *CCNB1* promoters contain CHR motifs^47^ required for MMB binding, increased expression of these cyclins mean that the G2/M transcription program was activated. Expression of this program generates a conflict between the S phase state and the G2/M state on the CDK1/2 spectrum. However, the transcription and translation cyclin A2/B1/CDK1 complexes requires time, explaining the delay in the generation of replication stress and DNA-PK/ATM activation.

The independent compensatory activities of DNA-PK and ATM following ATR inhibition also provide insight into ATM-proficient cells in which DNA-PK is reported to be the dominant compensator, including U2OS cells^23^, by suggesting that ATR inhibition induces an aberrant DNA replication structure during S phase that needs to be resolved prior to progressing onto G2. While DNA-PK and ATM classically are activated during DNA damage repair by double stranded breaks, KU70/KU80 complexes, which recruit and activate DNA-PK, can bind to stalled replication forks^48, 49^ while ATM drives the response to broken replication forks^10^, indicating that these kinases are activated by different aberrant DNA replication structures that appear in a sequential manner similar to the cell cycle checkpoints that were observed in this study. Alterations affecting telomere maintenance may also modulate DNA-PK activity independent of ATR, as 3 of 4 patients with tumors harboring *ATRX* mutations whose samples we assessed for pDNA-PK, had elevated basal levels. U2OS cells are also ATRX-deficient and consequently utilize alternative lengthening of telomeres (ALT), requiring HR machinery to replicate their telomeres^50^. Following ATR inhibition, DNA-PK activity during S phase may be critical to prevent competition between replicating telomeres and disassembling replication forks for access to the HR machinery necessary for the completion of genome duplication. Further investigation into whether oncogenic alterations, such as *ATRX* mutation, that may induce aberrant replication structures at telomeres to activate DNA-PK may identify additional patient cohorts that would benefit from combined ATRi + DNA-PK therapy.

These concepts will be tested in an ongoing Phase 1 study combining tuvusertib and peposertib (NCT05687136). Once the recommended Phase 2 dose of the combination is established, expansion cohorts are planned, including a group with *ATM*-mutant cancers, in whom ATM deficiency will be confirmed immunohistochemically, with the goal of realizing synthetic lethality that has not been adequately achieved with ATR inhibitor therapy alone.

## Materials and Methods

### Cell culture and drugs

SNU719 (CVCL_5086) and SNU1 (CVCL_0099) cells were originally purchased from ATCC and were propagated in RPMI 1640 medium (Life Technologies, Grand Island, NY) supplemented with 10% fetal bovine serum and 1% Pen/Strep (Gibco) at 37°C in 5% CO_2_. 500,000-700,000 cells were plated on 100 mm plates for treatment the next day and collection at the denoted time points post treatment. ATM was knocked out in SNU719 cells using a previously generated construct in lentiCRISPRv2 backbone (RRID:Addgene_52961) with sgRNAs targeting ATM^51^.

The following drugs were used in this study: tuvusertib (M1774; Merck KGaA, Darmstadt, Germany; CAS No. 1613200-51-3), peposertib (M3814; Merck KGaA, Darmstadt, Germany; CAS No. 1637542-33-6), lartesertib (M4076; Merck KGaA, Damstadt, Germany; CAS No. 2495096), gartisertib (M4344; Merck KGaA, Damstadt, Germany; CAS No. 1613191-99-3); RO3306 (Selleckchem; CAS No. 872573-93-8), CDK2-IN-4 (MedChemExpress HY-117535; CAS No. 2079895-42-2), KU55933 (MedChemExpress HY-12016; CAS No. 587871-26-9), nocodazole (Millapore-Sigma M1404; CAS No. 31430-18-9). For *in vitro* experiments, drugs were reconstituted and diluted in DMSO, while for *in vivo* experiments, the denoted formulations were used.

### Western blotting

Whole-cell extracts were generated by incubating cell pellets in RIPA Buffer (Boston BioProducts) with Protease Cocktail Set I (Millipore) and Phosphatase Cocktail Set I (Millipore) and cleared by centrifugation. Relative protein levels in lysates were normalized by Pierce BCA assay (Thermo Scientific) and samples were boiled in 6x SDS buffer for Tris-glycine gels or 4x LDS Sample Buffer and 10x NuPAGE Sample Reducing Agent (Invitrogen) for Bis-Tris gels. Samples were then loaded on Novex 4-20% Tris-glycine (Invitrogen) or NuPage 4-12% Bis-Tris gels (Invitrogen) for electrophoresis, followed by a wet transfer to nitrocellulose membrane (Bio-Rad). Membranes were blocked in 5% nonfat milk (Boston BioProducts) for 1 h at room temperature and incubated in primary antibody overnight at 4°C with dilutions listed in **Supplemental Table S1**. Membranes were incubated in secondary antibodies, either anti-Rabbit-HRP conjugate (Bethyl Laboratories A120-101P; RRID:AB_67264) or anti-mouse-HRP conjugate (Bethyl Laboratories A90-116P; RRID:AB_67183), diluted 1:5000 in 5% nonfat milk, for 1 h at room temperature. Bands were visualized using Super Signal West Pico (Thermo Scientific) or Immobilon Western Chemiluminescent (Millipore) HRP substrate on the G-box imaging system (Syngene) or ImageQuant LAS-4000 Mini (GE Healthcare). When reprobing, membranes were incubated twice in Restore Plus Stripping Buffer (Thermo Scientific) for 10 min at room temp before block and reprobing with primary antibodies as described above.

### Flow cytometry

Samples with BrdU or EdU incorporation were pulsed with 10 μM for 30 min prior to being collected with trypsin. Trypsinized cells were washed with PBS and fixed in 4% formaldehyde in PBS for 15 min at room temperature. Cells were then washed three times with 1% BSA in PBS and permeabilized in 70% ethanol at -20°C for 30 min to overnight. After three washes with 1% BSA in PBS, if the samples were pusled with BrdU, 500,000 cells were treated with DNase I (Sigma-Aldrich) for 1 hour at 37C or incorporated EdU was labeled with a CLICK reaction (2 mM CuSO4, 100 μM THPTA, 100 mM sodium ascorbate, and 2 μM Calfluor 647 Azide in PBS) by rotating for 30 min at room temperature. Following any DNase I treatment or click reaction, samples were then washed thrice with 1% BSA in PBS and incubated with the primary antibodies in 1% BSA in PBS overnight at 4⁰C as listed in **Supplemental Table S2**. Samples were washed three times in 1% BSA in PBS following primary antibody incubation and incubated for 1 hour at room temperature with secondary antibody as listed in **Supplemental Table S2**. If samples were pulsed with BrdU, samples were incubated overnight with anti-BrdU-APC conjugate following the completion of indirect staining and 3 washes with 1% BSA in PBS. Otherwise, after three washes 1% BSA in PBS washes, DNA staining was then conducted by incubating samples in darkness and at room temperature for 1 h in a 1 μg/ml DAPI, 100 ng/ml RNase A staining solution. Samples were analyzed on Fortessa analyzers (BD Biosciences). A negative control, where an EdU/BrdU pulse and primary antibodies were omitted but otherwise normally processed, was used to establish negative populations and perform background subtraction on MFIs.

For the chromatin flow cytometry, trypsinized cells were incubated on ice in 750 μl of Buffer A (300 mM sucrose, 100 mM NaCl, 3 mM MgCl_2_,10 mM PIPES pH 7.0, 0.5% Triton x-100, Protease Cocktail Set I, and Phosphatase Cocktail Set I) for 5 min. 250 μl of Buffer B (300 mM sucrose,100 mM NaCl, 3 mM MgCl_2_,10 mM PIPES pH 7.0, 0.5% Triton x-100, 10% formaldehyde, Phosphatase Cocktail Set I, and Phosphatase Cocktail Set I) was immediately added, and samples were incubated for an additional 15 min on ice. Cells were then pelleted (1000×g, 5 min, 4°C) and washed with 1% BSA in PBS 3 times. Otherwise, the CLICK reaction, primary and secondary as well as DAPI staining were conducted as described above and listed in **Supplemental Table 2**.

### Colony formation assays

For colony formation assays, 3,000 cells/well were plated in 6-well plates along with graded concentrations of drugs and cultured for 10–12 days for colony growth. Colonies were then fixed with methanol for 10 minutes at −20°C and stained with crystal violet. Plates were imaged with ImageQuant Las4000 (GE Healthcare) and quantification of the cell growth area was performed with ImageJ.

### Cell Viability Assays

500 cells in 100 µL of media were plated in each well of a 96-well plate and allowed to adhere overnight before the addition of drugged media resulting in the denoted final concentrations. Cells were treated with drug for six days and then cell viability was assessed using the CellTiter-Glo assay (Promega), according to the manufacturer’s instructions. Relative cell viability was normalized to the DMSO-treated control.

### Immunohistochemistry

For the staining of ATM expression in the PDX models, three-to-four-micron sections of formaldehyde-fixed paraffin-embedded (FFPE) tissues were mounted on hydrophilic adhesion slides and deparaffinized and stained using the Discovery® XT (Ventana Medical Systems, Inc., Tucson, USA; RRID:SCR_018643). A cell -line microarray using ATM high, medium and low expressing control cell lines served as run control.

For epitope retrieval, the slides were treated in Tris-EDTA buffer at pH 8 for 8 minutes at 96°C, followed by 28 minutes at 100°C. To quench endogenous peroxidase activity, the tissues were exposed to a 3% hydrogen peroxide solution (part of detection kit, Ventana, 760-4311) for 3 minutes at 37°C. The tissue sections were then incubated with the primary antibody for 32 minutes at 37°C. The primary antibodies used were rabbit anti-ATM clone Y170 (Abcam ab32420; RRID: AB_725574) or a rabbit isotype control (DA1E) mAb IgG XP (Cell Signaling Technologies 3900; RRID:AB_1550038). The binding of the primary antibody was detected using the OmniMap detection kit (Ventana) and developed with ChromoMap DAB (3′,3′-Diaminobenzidine; Ventana). Sections were briefly counterstained with hematoxylin (Ventana). After washing the slides in tap water, they were dehydrated and coverslipped in Entellan® Neu (VWR, Germany).

Finally, the slides were scanned using the NanoZoomer S210 (Hamamatsu; RRID: SCR_023760) and analyzed with HALO software (Indica Labs, v3.1; RRID:SCR_018350). Antibody staining (in arbitrary units) was quantified in viable tumor tissue regions using the area quantification module. In short, blue and brown staining are unmixed by color deconvolution and optical densities quantified for the brown staining. Antibody staining is then calculated by multiplying the percentage of viable tumor tissue with brown staining above background with the average optical density.

For the staining of pDNA-PK levels in patient samples from NCT03718091^21^, formalin-fixed core biopsy samples collected pretreatment and on-treatment within 30 hours of the fifth dose of berzosertib administration were dehydrated and embedded in paraffin. Sections (5 µm) were stained with pDNA-PK antibodies (Cell Signaling Technologies 68716S; RRID: AB_2939025) on the Leica BOND (RRID: SCR_025548) automated staining platform using recommended reagents. Stained slides were scanned and analyzed on the Aperio image analysis platform (RRID: SCR_020993) using previously determined cut-offs for staining intensity, classified as negative = 0, weak = 1, moderate = 2, and strong =3. H-scores (range 0–300) were estimated using the formula: H-score = [(3 X % nuclei with intensity 3) + 2 X % nuclei with intensity 2) + (% nuclei with intensity 1)]. Fold changes (FC) in H-scores were calculated using the formula FC (log_2_) = log2 (H-score on-treatment biospy) – log2 (H-score pretreatment biopsy).

### *In vivo* xenograft studies with SNU719 models

Experimental procedures involving the *in vivo* xenograft studies with SNU719 and SNU719-ATM-KO cells were conducted at Dana-Farber Cancer Institute (DFCI) with approval from the Institutional Animal Care and Use Committee (IACUC) and in compliance with AAALAC accreditation standards. All animal work followed the ethical guidelines established by DFCI’s IACUC. 7-week-old female NCr nude mice (Taconic Biosciences, NY) were used for the xenograft studies. A total of 5 × 10⁶ SNU719 or SNU719-ATM-KO cells were suspended in 100 µL of a 1:1 Matrigel/PBS mixture and injected subcutaneously in the right flank of mice. Animals were randomized into treatment groups when tumors reached an average tumor volume of 161.2 mm^3^ for the SNU719 tumors (n=5-8 mice/group) and 141.2 mm^3^ for the SNU719-ATM-KO tumors (n=6-7 mice/group). Mice were treated with vehicle control, peposertib (DNA-PKi) (100 mg/kg, twice daily, orally), gartisertib (ATRi-3) (10 mg/kg, daily, orally) or the combination of peposertib and gartisertib. Tumor growth was monitored by caliper measurements. Peposertib was formulated in 0.5% Methocel/0.25% Tween 20 in 300 mM citrate buffer pH 3.5. Gartisertib was formulated in 15% Captisol in water: 15% Captisol in 0.1M hydrochloric acid (95.04 : 4.96).

### PDX efficacy studies

In vivo efficacy studies were conducted using subcutaneous human patient-derived xenograft (PDX) models. The REX037 PDX model was established at ChemPartner, following Institutional Animal Care and Use Committee (IACUC) guidelines and standards of the Association for Assessment and Accreditation of Laboratory Animal Care (AAALAC). NU/NU mice were used, with tumor growth monitored biweekly. Mice with mean tumor volumes of 100-250 mm³ were randomized into treatment groups to ensure comparable mean tumor volumes across groups. The LXFA_2184 and LXFA_2155 PDX models were studied at Charles River Laboratories, adhering to international and national guidelines, with approval from the regional council Committee on the Ethics of Animal Experiments. NMRI nu/nu mice were monitored until tumors reached 50-250 mm³, and were assigned to vehicle or treatment groups for uniformity. The GAPF158 and CTC15008 PDX models were established at LIDE Shanghai Biotech, in compliance with LIDE IACUC. Tumors in nu/nu mice were measured, and when volumes reached 100-300 mm³, mice were divided into treatment groups.

For all studies, tumor volume (TV) was calculated using the formula TV=(l×w^2^) × 0.5, and body weight was recorded biweekly. Peposertib (DNA-PKi) was formulated in 0.5% Methocel, 0.25% Tween 20, and 300 mmol/L sodium citrate buffer (pH 2.5) for oral administration. Tuvusertib (ATRi) was prepared in 15% Captisol in water, adjusted to pH 4.75, and administered orally at specified doses.

### Quantification and statistical analysis

ImageJ was used to invert, adjust brightness, and crop immunoblot exposures for assembly of figures in Microsoft Powerpoint. Flow cytometry data was analyzed using FlowJo. GraphPad Prism was used to generate graphs and conduct statistical analyses for flow cytometry, immunohistochemistry, viability, and tumor growth data. All data are represented as mean ± SEM unless indicated otherwise. Significance was determine using one-way or two-way ANOVA with One-way or Two-way ANOVA with Tukey correction.

## Data Availability

The data supporting the findings of this study will be included in this article and its associated files. Source data will be provided with this paper.

## Acknowledgments

This work was supported by the Ludwig Center at Harvard (G.I.S.), a Sponsored Research Agreement from EMD Serono (CrossRef Funder ID: 10.13039/100004755; A.D.D., G.I.S.), as well as the Dana-Farber/Harvard Cancer Center (DF/HCC) Specialized Program of Research Excellence (SPORE) in Gastrointestinal Cancer, NIH Grant P50 CA127003 (A.D.D., G.I.S.) and the DF/HCC SPORE in Ovarian Cancer, NIH Grant P50 CA240243 (A.D.D., G.I.S.).

## Authors’ Contributions

**T.B. Branigan:** Conceptualization, investigation, data curation, formal analysis, methodology, supervision, writing–original draft, writing–review and editing. **R. Alharthi:** Investigation, data curation, writing–review and editing. **B.S. Kochupurakkal:** Investigation, methodology, writing– review and editing. **J.Hao:** Investigation, writing–review and editing. **S. Revia:** Investigation, data curation, writing–review and editing. **J. Albers:** Investigation, data curation, writing–review and editing. **M.-Y. Cai:** Investigation, writing–review and editing. **Y. Jiao:** Investigation, writing–review and editing. **K. McQueen:** Investigation, writing–review and editing. **S. Mukkavalli:** Project administration, writing–review and editing. **I.A. Roundtree:** Investigation, provision of patient samples, writing–review and editing. **G.M. Cote:** Investigation, provision of patient samples, writing–review and editing. **P.C. Gokhale:** Investigation, supervision, data curation, formal analysis, writing–review and editing. **K. Parmar:** Investigation, supervision, data curation, formal analysis, project administration, writing–review and editing. **A.D. D’Andrea:** Supervision, funding acquisition, writing–review and editing. **A .Zimmerman:** Investigation, supervision, data curation, formal analysis, writing–review and editing. **G.I. Shapiro:** Conceptualization, supervision, project administration, acquisition of funding, formal analysis, writing–original draft, writing–review and editing.

