## Supplementary Information for "DNA-PK enforces the S/G2 checkpoint in ATM-deficient cells to limit ATR inhibitor efficacy"

Timothy B. Branigan^1,2,3^, Roshen Alharthi^1^, Bose S. Kochupurakkal^1,4^, Jie Hao^1,2,3^, Steffie Revia^5^, Joachim Albers^5^, Mu-Yan Cai^3,6^, Yuqing Jiao^1^, Kelsey McQueen^4,6^, Sirisha Mukkavalli^4,6^, Ian A. Roundtree^3,7^, Gregory M. Cote^3,7^, Prafulla C. Gokhale^8^, Kalindi Parmar^3,4,6^, Alan D. D’Andrea^3,4,6,9^, Astrid Zimmermann^5^,

Geoffrey I. Shapiro^1,2,3,4,9^

^1^Department of Medical Oncology, Dana-Farber Cancer Institute, Boston, Massachusetts

^2^Department of Medicine, Brigham and Women’s Hospital, Boston, Massachusetts

^3^Harvard Medical School, Boston, Massachusetts

^4^Center for DNA Damage and DNA Repair, Dana-Farber Cancer Institute, Boston, Massachusetts

^5^Research Unit Oncology, Healthcare Business of Merck KGaA, Darmstadt, Germany

^6^Department of Radiation Oncology, Dana-Farber Cancer Institute, Boston, Massachusetts
^7^ Mass General Brigham Cancer Institute, Boston, Massachusetts, USA

^8^Belfer Center for Applied Cancer Science, Dana-Farber Cancer Institute, Boston, Massachusetts

^9^Ludwig Center at Harvard, Boston, Massachusetts

**Running Title:** Distinct compensation by DNA-PK and ATM after ATR inhibition

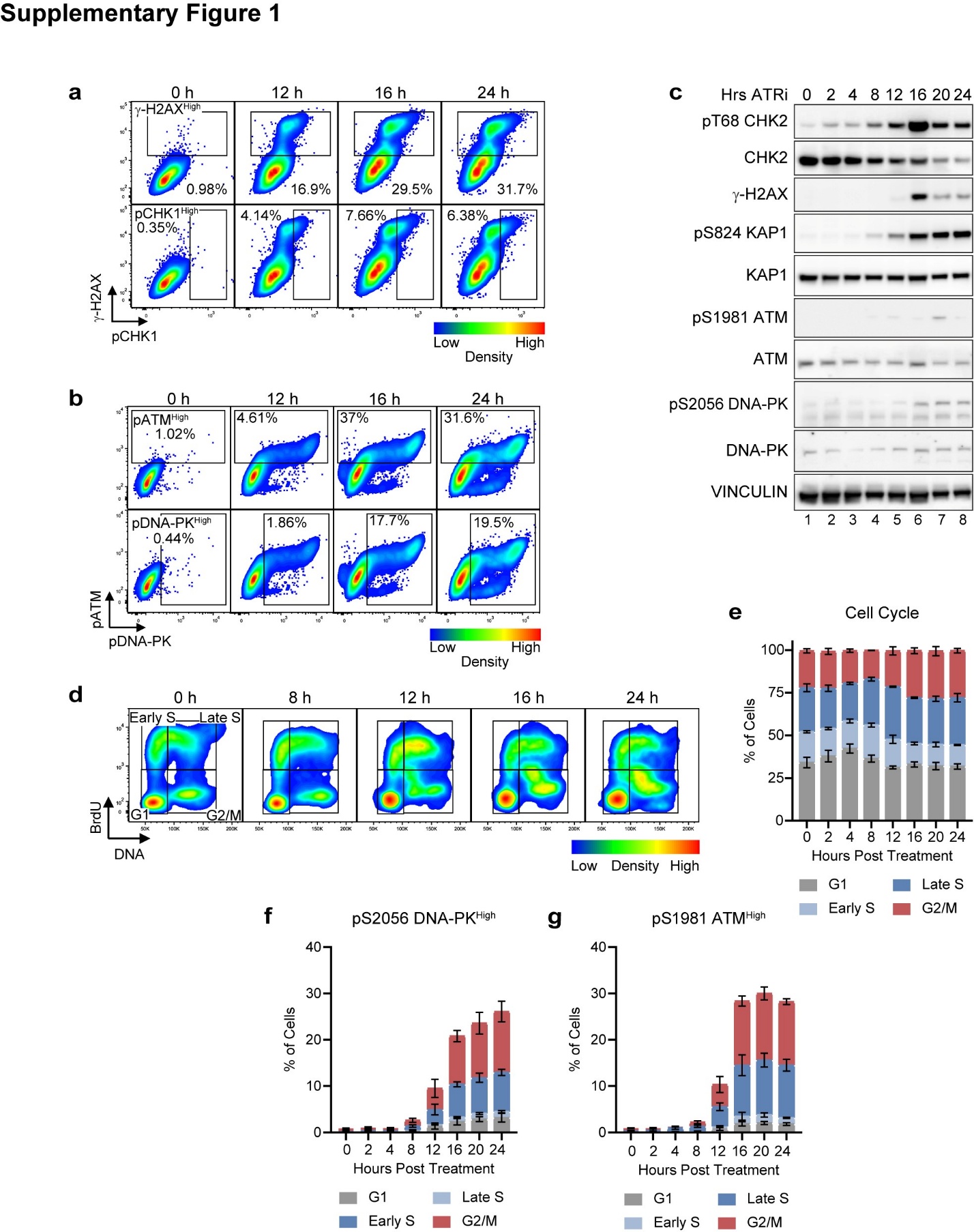

**Supplementary Figure 1. Cell cycle-dependent activation of DNA-PK and ATM following ATR inhibition.** Activation of DNA-PK and ATM at an S/G2 checkpoint following tuvusertib (ATRi) treatment. SNU719 cells were treated with 500 nM ATRi for the denoted time points and either pulsed with BrdU and analyzed by flow cytometry after staining for γ-H2AX, pCHK1, pDNA-PK, pATM, and DNA content, N=3, or harvested to generate whole cell extracts and probed. **a** Gates for γ-H2AX^High^ (Top) and pCHK1^High^ (bottom) at 0, 12, 16, 24 hours post ATRi treatment. **b** Gates for pATM^High^ (Top) and pDNA-PK^High^ (bottom) at 0, 12, 16, 24 hours post ATRi treatment. **c** Immunoblot of CHK2 activation and induction of pKAP1, γ-H2AX^High^, pATM, pDNA-PK following ATRi treatment. **d** Cell cycle distribution at 0, 8, 12, 16, 24 hours post ATRi treatment **e** Cell cycle population distribution of samples following ATRi **f** % cells that are pDNA-PK^High^ colored based on cell cycle population **g** % cells that are pATM^High^ colored based on cell cycle population.

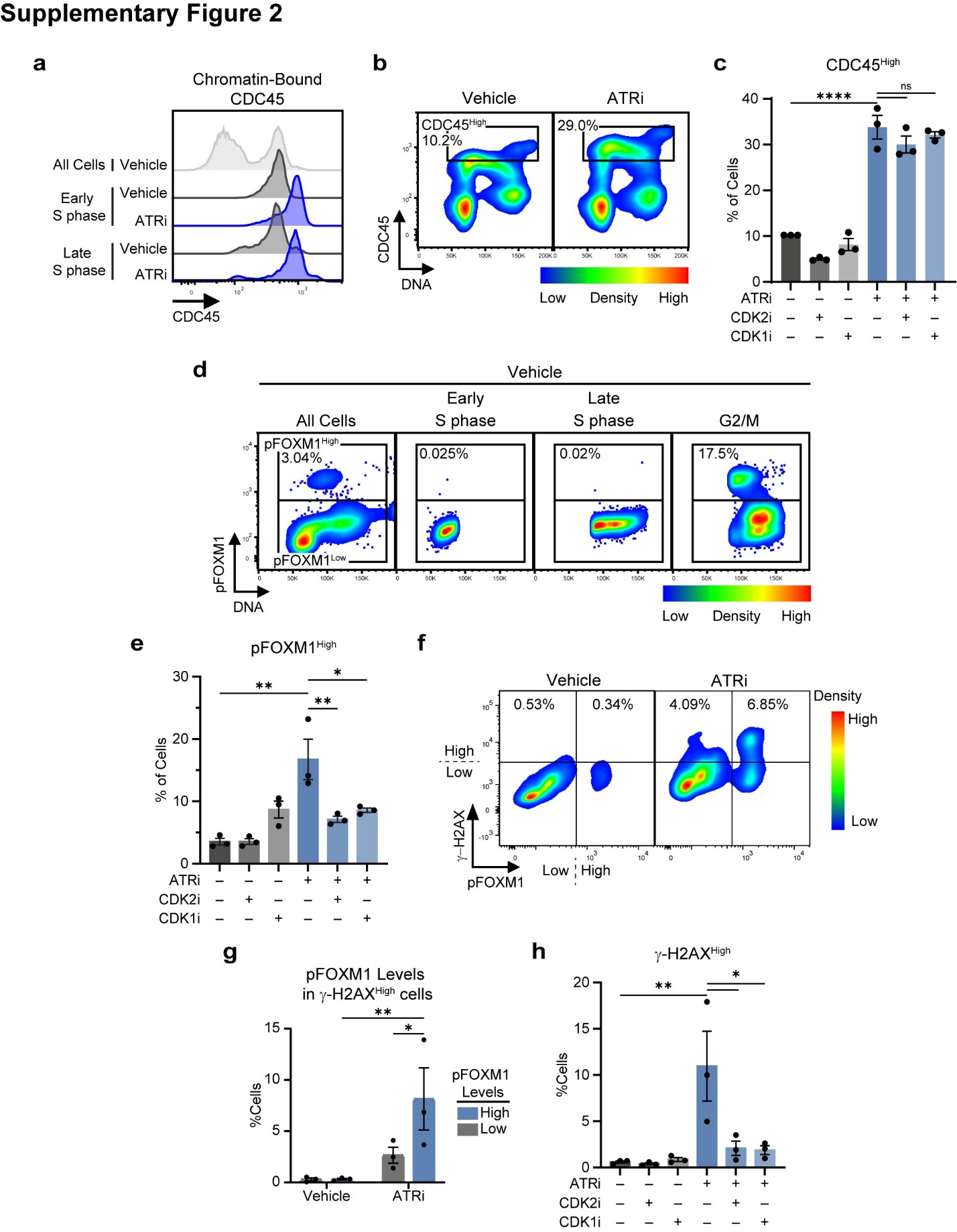

**Supplementary Figure 2. Sufficient CDKi to block DNA damage follow ATRi prevent premature S to G2 transition but not high origin firing.**

**a-c** CDKi inhibition sufficient to block DNA-PK and ATM activation fail to block origin firing following ATRi (tuvusertib) treatment, related to **Fig. 1j**. SNU719 cells were treated with 500 nM ATRi ±10 μM CDK2i (CDK2-IN-4) and/or 2.5 μM CDK1i (RO3306) for 16 hours. Samples were pulsed with EdU and analyzed by chromatin flow cytometry following staining for chromatin-bound CDC45, EdU, and DNA content, N = 3. **a** Modal histogram comparing levels of chromatin-bound CDC45 in Early and Late S phase in vehicle and ATRi treated samples. **b** Levels of chromatin-bound CDC45 compared with DNA content in the vehicle and ATRi treated samples with the CDC45^High^ gate denoting origin firing above the ~90^th^ percentile of the vehicle samples. **c** % of cells that are CDC45^High^. **d-f** ATRi induces pFOXM1 in Late S phase to compromise the S/G2 checkpoint, related to **Fig. 1k-m**. SNU719 cells were treated as above. Samples were pulsed with BrdU and analyzed by flow cytometry following staining for γ-H2AX, pT600 FOXM1, BrdU, and DNA content, N = 3. **d** Levels of pFOXM1 compared with DNA content in the overall sample (left), Early S phase (Middle Left), Late S phase (Middle Right), and G2/M (right). % denotes the % of pFOXM1^High^ cells within the respective population. **e** % of cells that are pFOXM1^High^ **f** Comparing high and low levels of pFOXM1 and γ-H2AX^High^ following ATRi. **g** % of cells that are γ-H2AX^High^ separated based on pFOXM1 Levels, **h** % of cells that are γ-H2AX^High^. * p<0.05. ** p<0.01. *** p<0.001. **** p<0.0001.

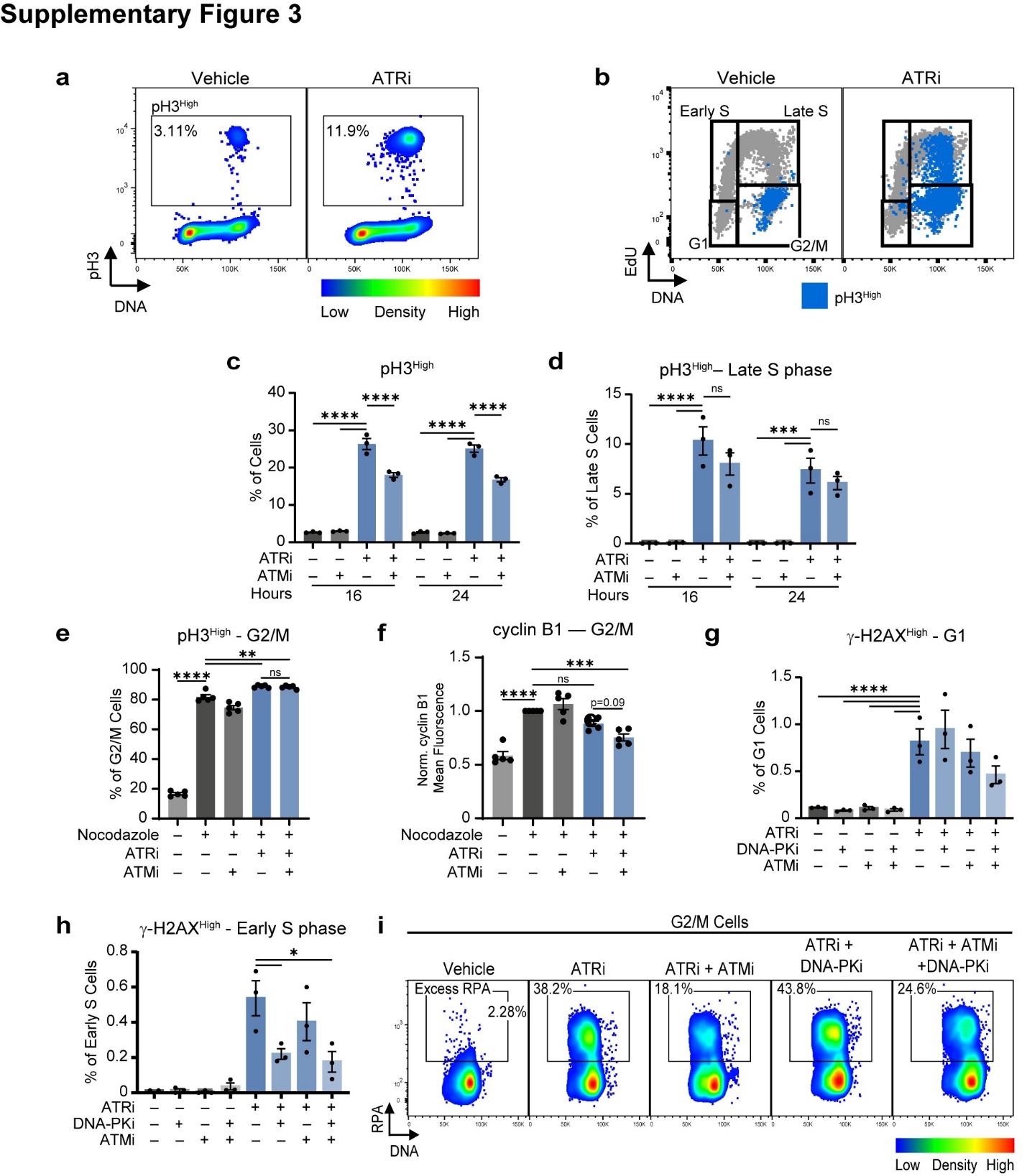

**Supplementary Figure 3. ATM enforces a G2/M checkpoint following ATR inhibition while DNA-PK inhibition detects DNA damage in Early S phase.** **a-h** Differing roles of DNA-PK and ATM in enforcing cell cycle checkpoints and detecting DNA damage, related to **Fig. 2c-h**. SNU719 cells were treated with 500 nM ATRi ± 2.5 μM DNA-PKi and/or 2.5 μM ATMi for 16 hours or, in the case of c-e, 16 and 24 hours. Levels of γ-H2AX, pH3, EdU incorporation, and DNA content were analyzed by flow cytometry, N=3. **a** Identification of a pH3^High^ population by comparing pH3 levels with DNA content. **b** pH3^High^ cells within the cell cycle. pH3^High^ cells in blue overlaid on to cell cycle distribution. **c** % of that are pH3^High^ at 16 and 24 hours post treatment following ATRi ± ATMi. **d** % of Late S phase cells that are pH3^High^ at 16 and 24 hours post treatment following ATRi ± ATMi.

**e** and **f** ATM maintains the G2/M state following ATR inhibition. SNU719 cells treated as above for 24 hours with 100 ng/ml nocodazole added at 12 hours. Levels of pH3, cyclin B, EdU incorporation, and DNA content were analyzed by flow cytometry, N=5. **e** % of G2/M cells that are pH3^High^. **f** mean fluorescence intensity of cyclin B1 in G2/M normalized to the nocodazole-treated vehicle. **g** % of G1 cells that are γ-H2AX^High^. **h** % of Early S phase cells that are γ-H2AX^High^. **i** Chromatin-bound RPA levels compared to DNA content in the G2/M population with gate denoting G2/M cells with excess RPA, related to **Fig. 2i**. SNU719 cells were treated with 500 nM ATRi ± 2.5 μM ATMi, and/or 2.5 μM DNA-PK for 16 hours. Chromatin flow cytometry was performed for chromatin-bound RPA levels, EdU incorporation, and DNA content. * p<0.05. ** p<0.01. *** p<0.001. **** p<0.0001.

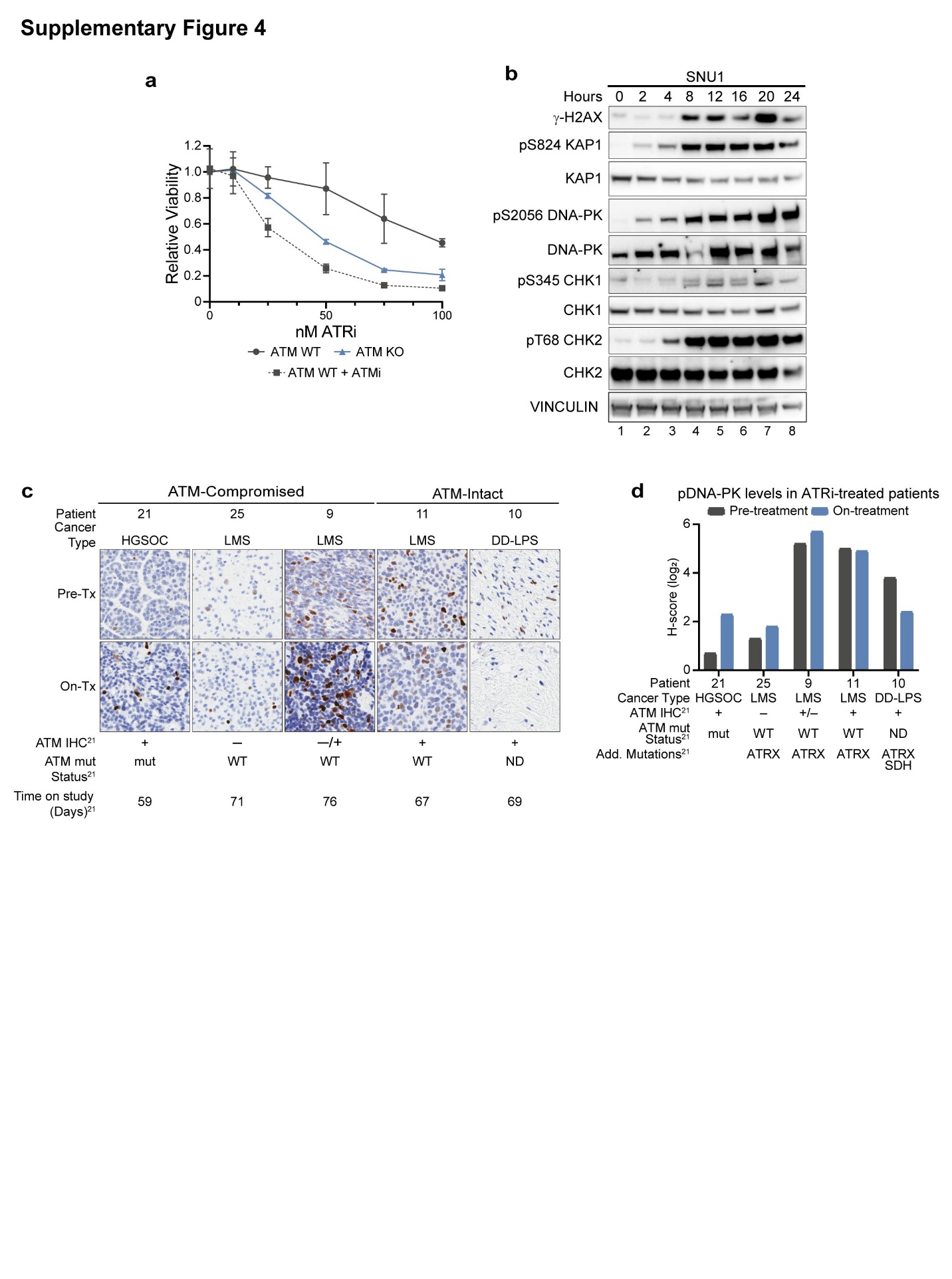

**Supplementary Figure 4. DNA-PK is activated in ATM-compromised systems after ATR inhibition. a** Cell viability assays of SNU719 ATM WT and KO cells treated with denoted concentrations of ATRi (tuvusertib) ± 400 nM ATMi, N=2. Mean ± SD. **b** Immunoblot of replication stress marker induction and activation of CHK1/2 following 500 nM ATRi in SNU1 cells. Whole cell extracts were generated and probed with denoted antibodies. **c** and **d**) Activation of DNA-PK in patients with ATM-compromised tumors following treatment with ATRi-2 (berzosertib) (NCT03718091). **c** Immunohistochemistry staining for pDNA-PK in pre-treatment and on-treatment patient samples. ATM status and time on study are indicated. **d** Quantification of pDNA-PK IHC staining in C) with ATM status and additional mutations listed.

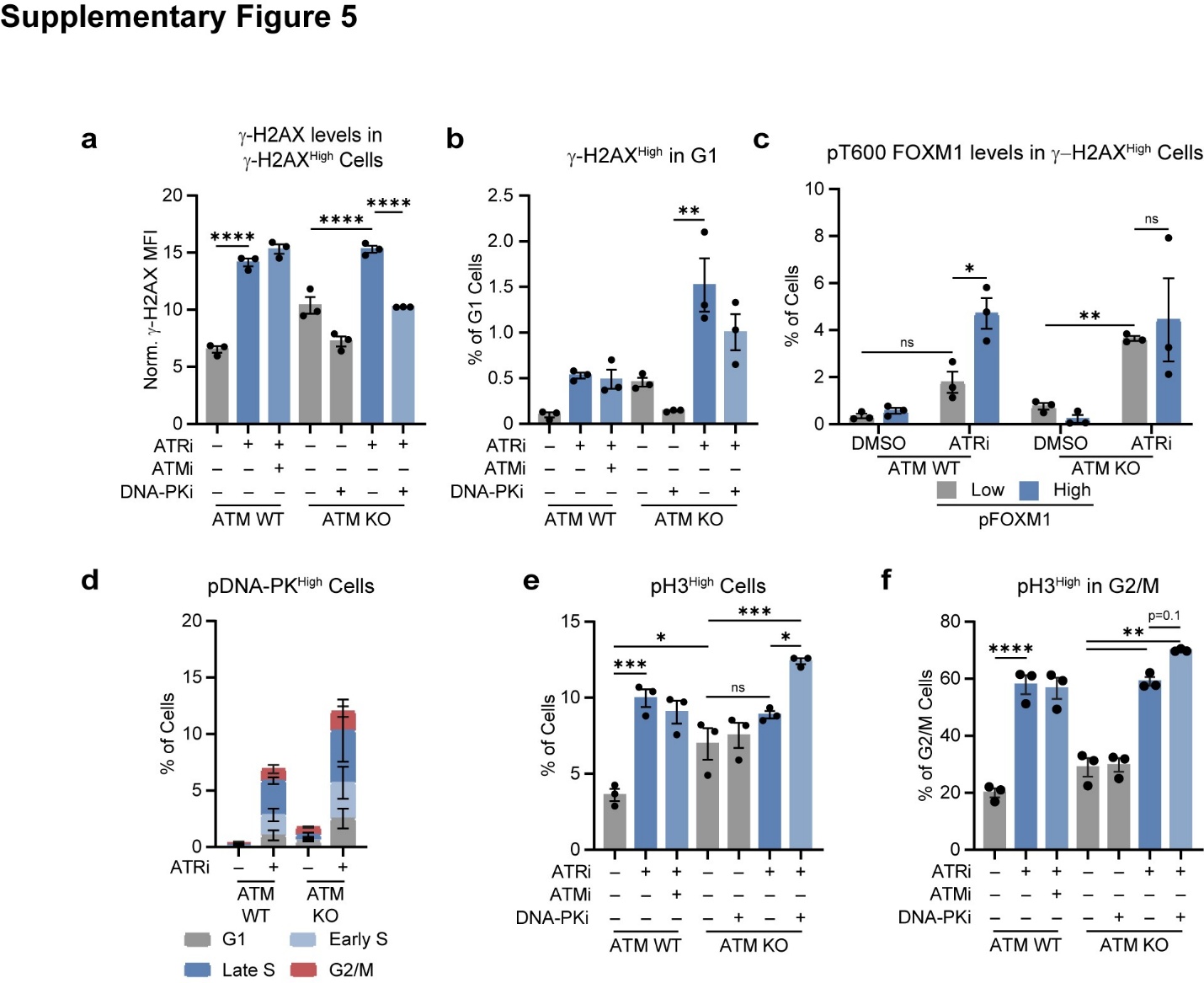

**Supplementary Figure 5. Cell cycle-dependent compensation by DNA-PK following ATR inhibition in ATM KO cells. a** and **b** DNA-PK detects DNA damage in ATM KO cells, related to **Fig. 3e and 3d**. ATM WT and ATM KO SNU719 cells were treated with 500 nM ATRi, 2.5 μM ATMi, and/or 2.5 μM DNA-PK as denoted for 16 hours. Levels of γ-H2AX, pH3, EdU incorporation, and DNA content were analyzed by flow cytometry, N=3. **a** Mean Fluorescence Intensity of γ-H2AX in γ-H2AX^High^ cells. Normalized to γ-H2AX MFI of DMSO sample. **b** % of G1 cells that are γ-H2AX^High^. **c** and **d** DNA damage is detected early in cells division following ATRi (tuvusertib) in ATM KO cells. ATM WT and KO SNU719 cells were treated with 500 nM ATRi for 16 hours and levels of γ-H2AX and pFOXM1 or pDNA-PK, BrdU incorporation, and DNA content were analyzed by flow cytometry, N=3. **c** % cells that are γ-H2AX^High^ separated based on pFOXM1 Levels, **d** % cells that are pDNA-PK^High^ colored based on cell cycle population. **e** and **f** DNA-PK detects DNA damage in S phase and reinforces the S/G2 checkpoint following ATRi in ATM deficient cells, related to **Fig. 3g-j**, (**a, b**), N=3. **e** % of cells that are pH3^High^, **f** % of G2/M cells that are pH3^High^. * p<0.05. ** p<0.01. *** p<0.001. **** p<0.0001.

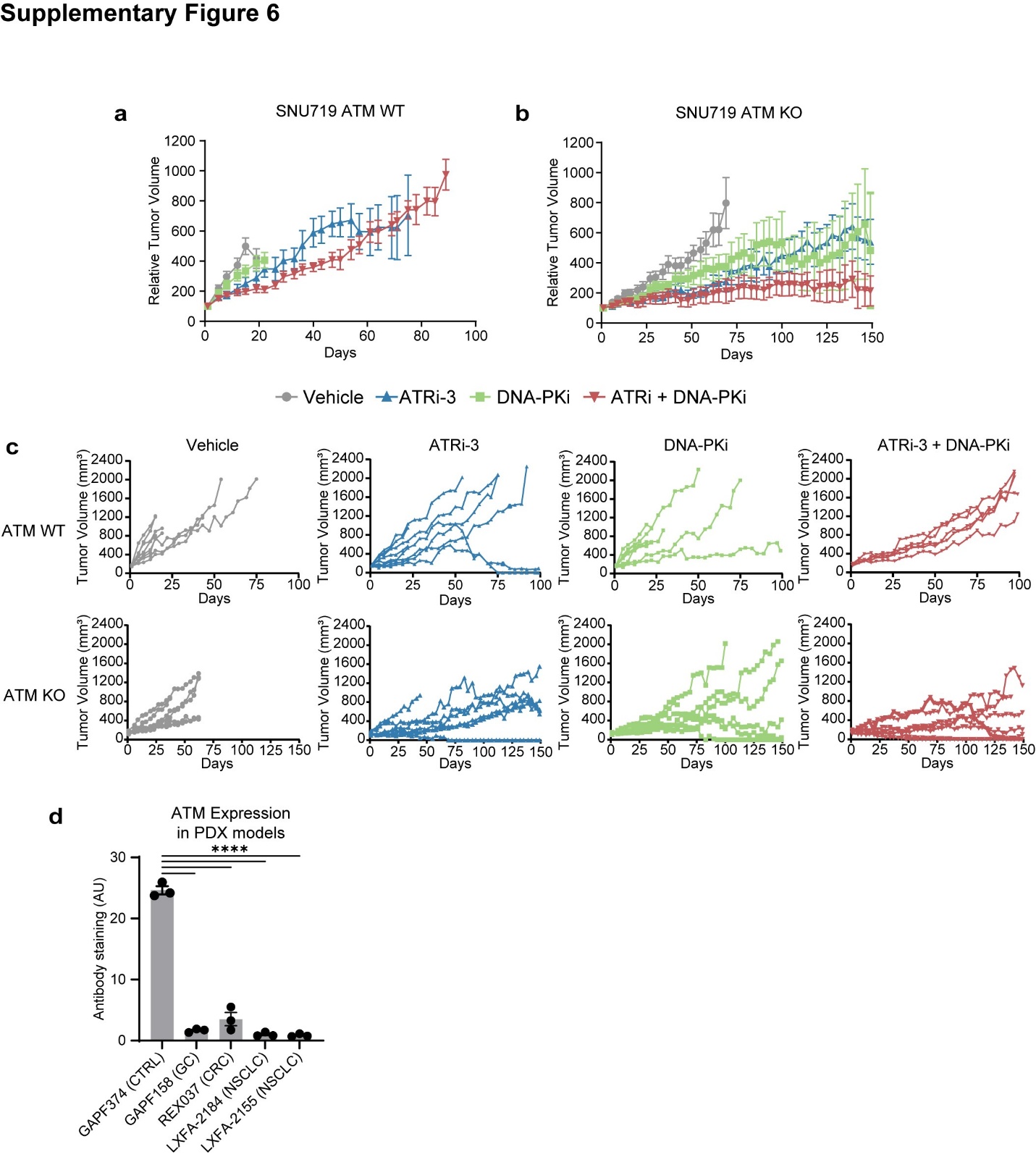

**Supplementary Figure 6. ATM-deficient tumors are sensitive to combined ATR + DNA-PK inhibitor therapy.** **a-c** DNA-PKi (peposertib; 100 mg/kg; BID) enhances ATRi-3 (gartesertib; 10 mg/kg; PO; QD) efficacy in SNU719 xenografts. **a** and **b** Relative tumor volumes of **a** ATM WT and **b** ATM KO SNU719 xenografts after ATRi-3 + DNA-PKi combination therapy at time points with ≥5 mice. **c** Tumor volumes in individual mice over time with ATM WT (top) and ATM KO (bottom) tumors treated with vehicle (left), ATRi-3 (center left), DNA-PKi (center right), ATRi-3 + DNA-PKi (right) **d** Quantification of ATM expression levels in the IHC samples of the PDX models in **Fig. 4e**. **** p<0.0001.

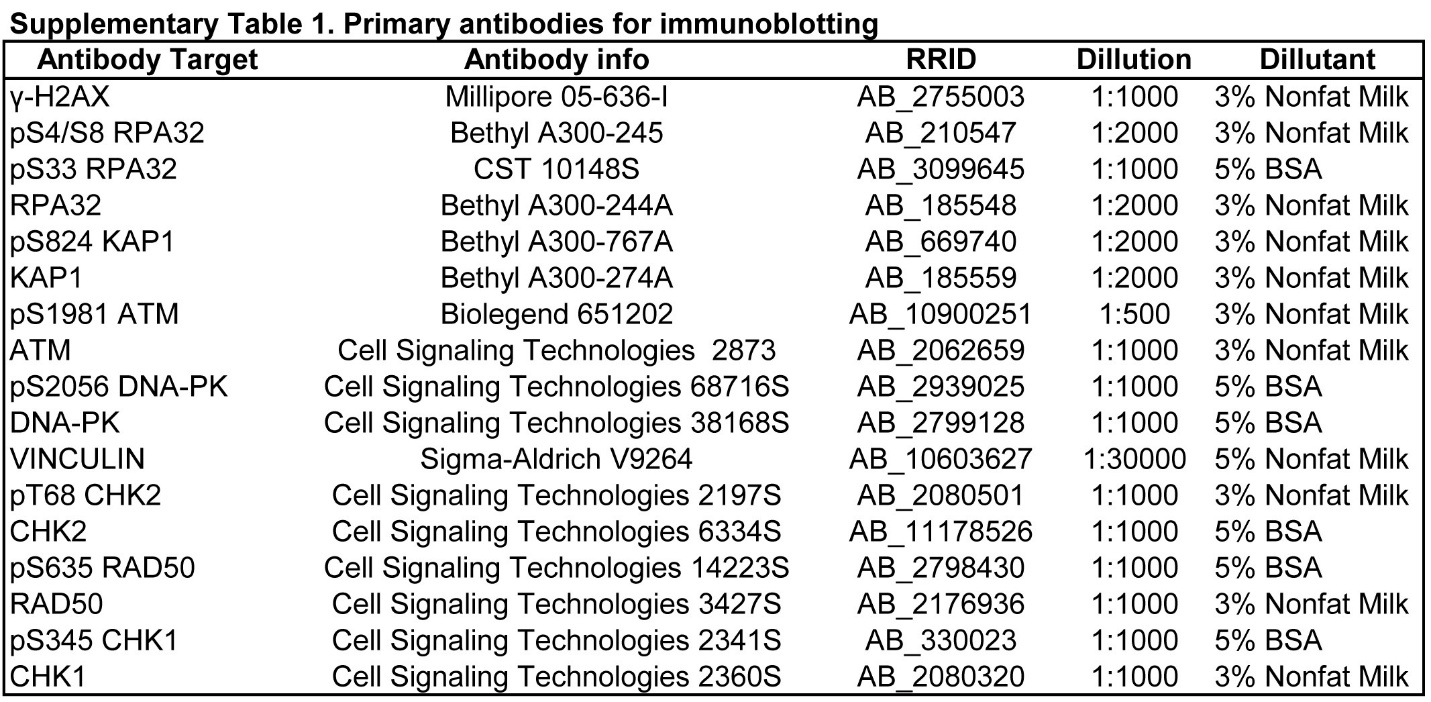

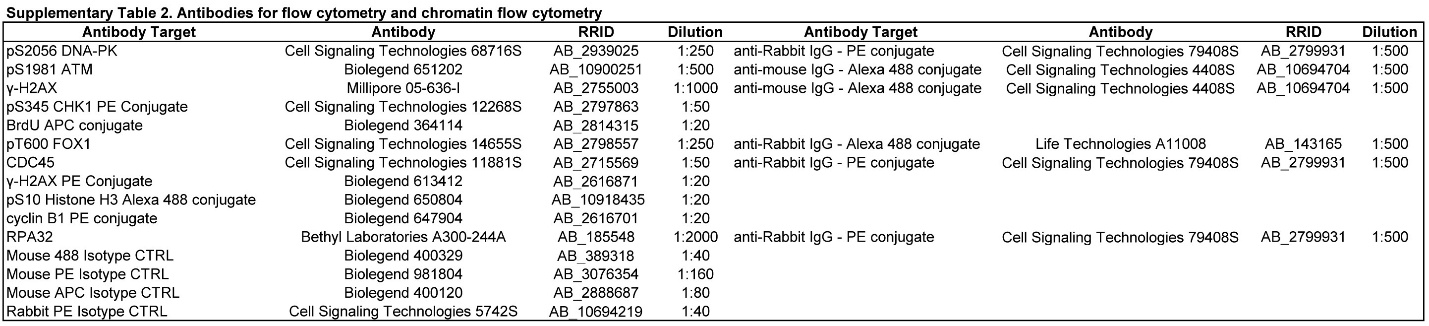
